# Prediction of DNA from context using neural networks

**DOI:** 10.1101/2021.07.28.454211

**Authors:** Christian Grønbæk, Yuhu Liang, Desmond Elliott, Anders Krogh

## Abstract

One way to better understand the structure in DNA is by learning to predict the sequence. Here, we train a model to predict the missing base at any given position, given its left and right flanking contexts.

Our best-performing model is a neural network that obtains an accuracy close to 54% on the human genome, which is 2% points better than modelling the data using a Markov model. In likelihood-ratio tests, we show that the neural network is significantly better than any of the alternative models by a large margin. We report on where the accuracy is obtained, observing first that the performance appears to be uniform over the chromosomes. The models perform best in repetitive sequences, as expected, although they are far from random performance in the more difficult coding sections, the proportions being ~ 70:40%. Exploring further the sources of the accuracy, Fourier transforming the predictions reveals weak but clear periodic signals. In the human genome the characteristic periods hint at connections to nucleosome positioning. To understand this we find similar periodic signals in GC/AT content in the human genome, which to the best of our knowledge have not been reported before.

On other large genomes similarly high accuracy is found, while lower predictive accuracy is observed on smaller genomes. Only in mouse did we see periodic signals in the same range as in human, though weaker and of different type. Interestingly, applying a model trained on the mouse genome to the human genome results in a performance far below that of the human model, except in the difficult coding regions.

Despite the clear outcomes of the likelihood ratio tests, there is currently a limited superiority of the neural network methods over the Markov model. We expect, however, that there is great potential for better modelling DNA using different neural network architectures.

## Introduction

We consider the question of how predictable chromosomal DNA is from context. That is: how well can one predict the base at any given position in the DNA, if the surrounding context of bases around the position are known, e.g. the five bases to the left and right of the position. The motivation for this approach is that the surrounding context exerts some kind of pressure on a missing base. Such pressure could stem from, or be connected with, structured parts of the genome.

In our parallel work [10], we use conditional probability models with contexts up to 14 nucleotides on side each (29 in total) around the central nucleotide. It is likely that larger contexts influence the probability of a nucleotide, because many structures in the genome, such as genes, nucleosomes, and transposable elements, are larger than 29 nucleotides. In the present study we use neural networks to model nucleotide contexts, which allows for larger contexts, and compare the performance of neural network-based models against the conditional probability models, which are forced to use contexts of smaller size as input, because the number of parameters otherwise explodes.

Concretely, we formulate the question in terms of estimating the probability of the four possible bases at a given position in the genome, conditioned on the knowledge of the bases in *flanks* of equal length to the left and to the right of the position [10]. Using common notation these probabilities can be written

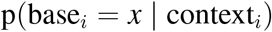

where base_*i*_ is one of the four bases, A, C, G and T, and the context_*i*_ is given by the *left flank* and *right flank*, which are the sequences of bases to the left and to the right of position *i*. Having modelled these probabilities, we let the model predict the missing base as the most probable assignment 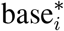:

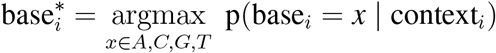

A measure of the prediction made by a model is its accuracy, which is the fraction of all positions at which the model assigns the true base.

Our baseline model [10], referred to as the *k* = 3 central model, uses contexts of size 6, i.e. three bases in the left and right flanks around the missing position. The ~12000 model parameters (three times the number of contexts of size 6) are estimated simply by counting for a given context the number of times each of the four letters occurs in the genome; the corresponding frequencies can then be taken to be the desired conditional probabilities. On the human genome, the *k* = 3 central model obtains an accuracy of ~38%. With flanks of size 5 or 7, i.e. for a *k* = 5 or *k* = 7 central model, the accuracy increases to ~45%, and ~49%, respectively, and the number of parameters rises to ~3 million, and ~0.8 billion, respectively. Of these ‘‘simple models” only a Markov model estimated on the left-hand contexts of size *k* = 14 broke through 50% accuracy (~ 52% using the left-hand flanks on both strands for the predictions); this model has the same number of parameters as the *k* = 7 central model.

Given that best performance of the Markov model reaches 52%, it is natural to consider whether improved accuracy can be achieved using bigger contexts, e.g. 50 or 100 bases. In the case of the statistical models, the frequencies used to estimate a central model over such large contexts result in statistics that are too sparse: the model parameters outnumber the input data points. To remedy this, in [10] we let already the *k* = 5 central model be interpolated on the *k* = 4 central model, and proceed in this way for k’s above five, and the same is done for the Markov model. We turn to neural networks language models to overcome this curse of dimensionality [3]. We construct our neural networks so they can process larger contexts than the previous models and yield a single prediction for the value of the missing base. For defining the larger architecture of the neural networks relevant to our task, an obvious source of inspiration is their use in natural language modelling tasks. This points to using recurrent neural networks with Long Short Term Memory hidden units (LSTM [7]) or Transformer networks ([17]). We have also carried out experiments using feed forward networks and convolutional networks ([6] [9]).

The best performing networks we have found consist of one or two convolutional layers for “word encoding” (tri- or quadro-nucleotides) followed by bidirectional LSTM layers. On the human genome, the best-performing model in terms of accuracy goes beyond 53 %, and averaging bi-directionally, as with the Markov model, increases performance by a further ~0.5 %. To formally compare the best neural network with the simpler models we conduct a likelihood-ratio test for not nested models [19].

We also train the same or similar models on genomes of other organisms, resulting in highly varying accuracy. One option here is to obtain a measure of genomic similarity, by applying a model trained on one organism’s genome to another organism’s genome. As an example we compare a) the predictions obtained by applying a model trained on mouse DNA to human DNA to b) the predictions obtained by applying a model trained on human DNA to its own DNA. The results are somewhat surprising: of the annotated matter we consider, the two models seem to be in fine agreement only in coding regions, while elsewhere there are large differences, in particular in repeats.

It is also of interest to look into where a model performs well. It is, for instance, to be expected that much of the accuracy obtained can be due to repetitive DNA sequences. We show how the accuracy varies over annotated matter, such as various types of repetitive sequences, coding regions and more. Using Fourier transformation, we also investigate whether the predictions of a model carry signs of periodic components. This depends on the considered organism: in the genomes considered here we find clear signals only in human, persistently shared across the chromosomes. On the scale of the highest amplitudes in the Fourier coefficients these signals are small. For mouse such signals are surprisingly much less clear, verging on non-existent; in zebrafish, yeast and fruit fly there are no consistent signs of this kind through the genomes.

For the human genome we found quite similar signals in GC/AT content (which to the best of our knowledge have not been reported before). The GC/AT periods are slightly different and the amplitudes also differ from those we find in the models’ predictions. The latter periods quite clearly suggest connection with DNA wrapping/condensation (nucleosome distance and length of linker DNA), underpinned by the similarity to the GC/AT content case. In the other genomes we found some vaguely similar structure in the GC/AT content only in mouse.

## Results

### Human genome

We trained three multilayer neural network models of convolutional-LSTM type (LSTM1, LSTM11, LSTM4) on the human reference genome hg38, and compared their accuracy when applied to the same genome. The training data consisted of about one third of the genome and the size of the validation set was one-fifth of that (for details on the networks and their training see Supplementary Methods). An additional model, LSTM4S, also of convolutional-LSTM type, was trained only on the odd-numbered chromosomes of hg38. The LSTM1 model uses flanks of size 200; LSTM11 is identical to LSTM1 but trained less; LSTM4 has almost the same architecture as these models, and LSTM4S is identical to LSTM1, but both use flanks of size 50. The reason for including LSTM11 was to investigate where the increase in accuracy from the extra training stems from, while LSTM4 and LSTM4S are included to shed light on the impact of the context size (though LSTM4S’s main role is to ascertain our training-testing approach, see Supplementary Methods).

#### Accuracy

Figure 1 shows the performance of the three models in terms of accuracy per autosomal chromosome (all values can be found in Table 1 in the Supplementary Tables and Plots). The LSTM1 shows invariably the best performance (overall ~ 53%), closely followed by LSTM4 and then LSTM11, the less trained version of LSTM1. The very same pattern is seen when the accuracy is grouped by annotation (Figure 2).

**Figure 1:**
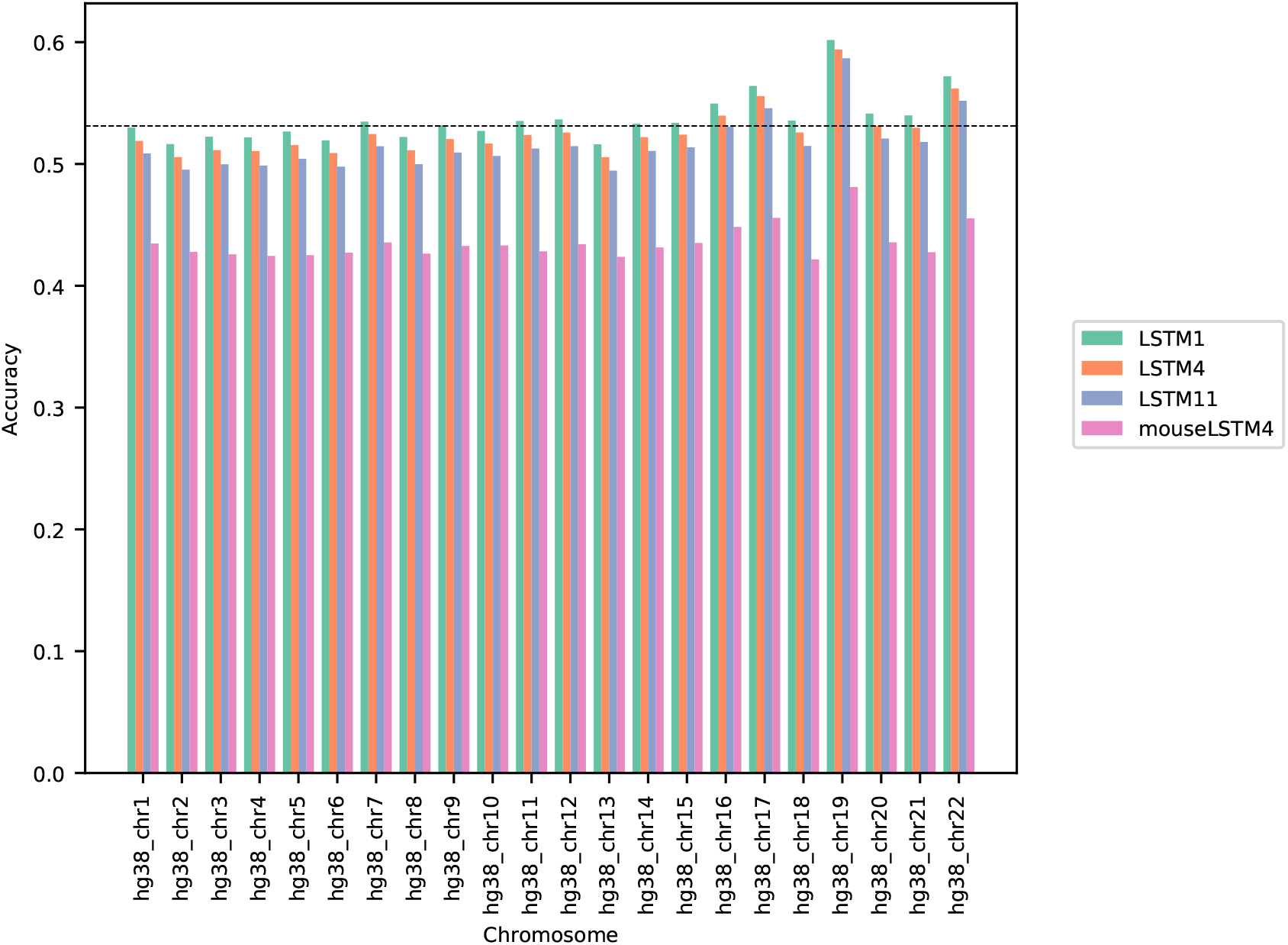
Model accuracy comparison across the human autosomal chromosomes. The named models are described in the text. The dashed line indicates the average performance of the LSTM1. All values can be found in Table 1 in the Supplementary Tables and Plots.

**Table 1:**
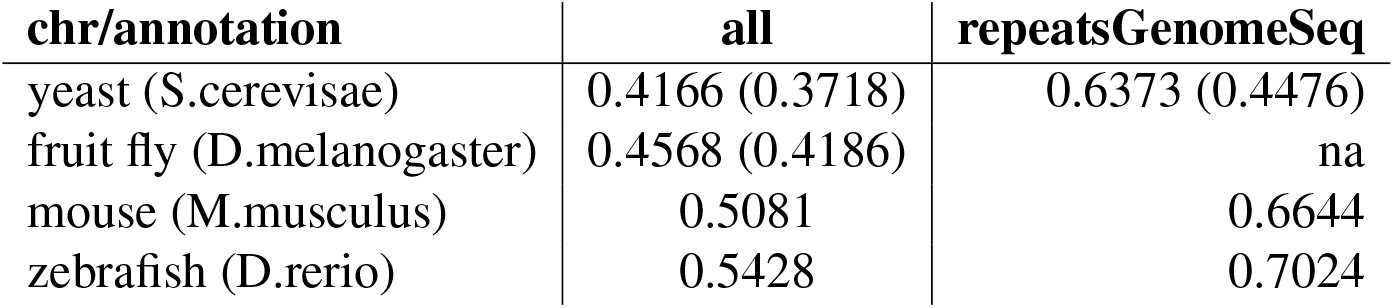
Overall accuracy for the LSTM4 trained on reference genomes of the named organisms. The numbers in parentheses are due to the early stage models for yeast (LSTM41) and fruit fly (LSTM41), see Supplementary Methods.

| chr/annotation | all | repeatsGenomeSeq |
| --- | --- | --- |
| yeast (S.cerevisae) | 0.4166 (0.3718) | 0.6373 (0.4476) |
| fruit fly (D.melanogaster) | 0.4568 (0.4186) | na |
| mouse (M.musculus) | 0.5081 | 0.6644 |
| zebrafish (D.rerio) | 0.5428 | 0.7024 |

**Figure 2:**
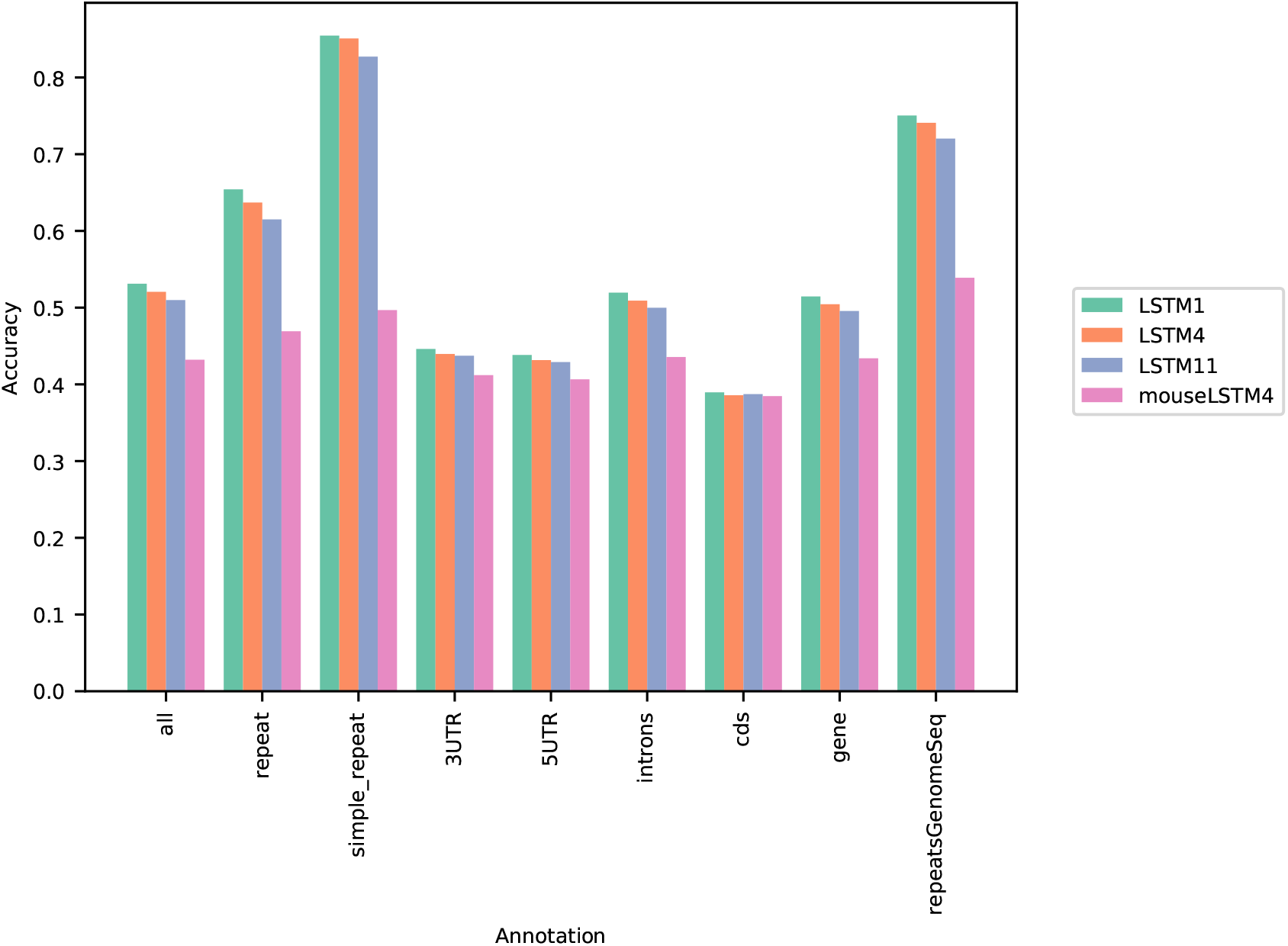
Four model accuracy comparison by annotation. All values can be found in Table 2 in the Supplementary Tables and Plots.

The differences between LSTM1 and LSTM4 suggest that there is additional information in the larger contexts. Except for a more rich “word-encoding” convolutional part, LSTM1 differs from LSTM4 only in the use of larger flanks, and the models have almost the same number of trainable parameters (LSTM1 2,227,582 vs. LSTM4 2,135,806). This is also supported by the results of LSTM4S, which are only marginally different from those of LSTM4 (see Supplementary tables and plots and Supplementary Methods for results of LSTM4S and comparisons to the other models); the differences in performance between LSTM1 and LSTM4S stem from the different flank sizes and the differences in their training sets (on the odd-numbered chromosomes LSTM4S has an advantage, but the superiority of LSTM1 seems largely uniform across the chromosomes). We also saw in experiments that shuffling the contexts while retaining some few central bases led to some loss in accuracy (data not included).

The differences between LSTM1 and LSTM11 show broadly that the increase in performance from LSTM11 to the (much) more trained LSTM1 is not ascribable to particular chromosomes or annotated parts of the genome: The increase is about the same across chromosomes and across annotations.

Figure 2 shows that as expected a large part of the models’ accuracy is obtained in the repeats, where they invariably perform much better than elsewhere. Whereas the total accuracy (‘all’) is slightly above 50%, it is about 65% in the repetitive sequences as defined by the repeat masking in the downloaded genome sequence [16]. The difficulties in predicting are most pronounced in coding regions, though the accuracy is still close to 40% there. All values can be found in Table 2 in the Supplementary Tables and Plots.

As can be seen in Figure 3, the accuracy level appears stable across the chromosomes (only data for LSTM1 are included). The largest variation is seen for simple repeats, and for (the GC rich) chromosome 19 there are some differences to the other chromosomes across the annotations. Also notable, the ‘gene’ category appears very close to the total, ‘all’. All values can be found in Table 3 in the Supplementary Tables and Plots.

**Figure 3:**
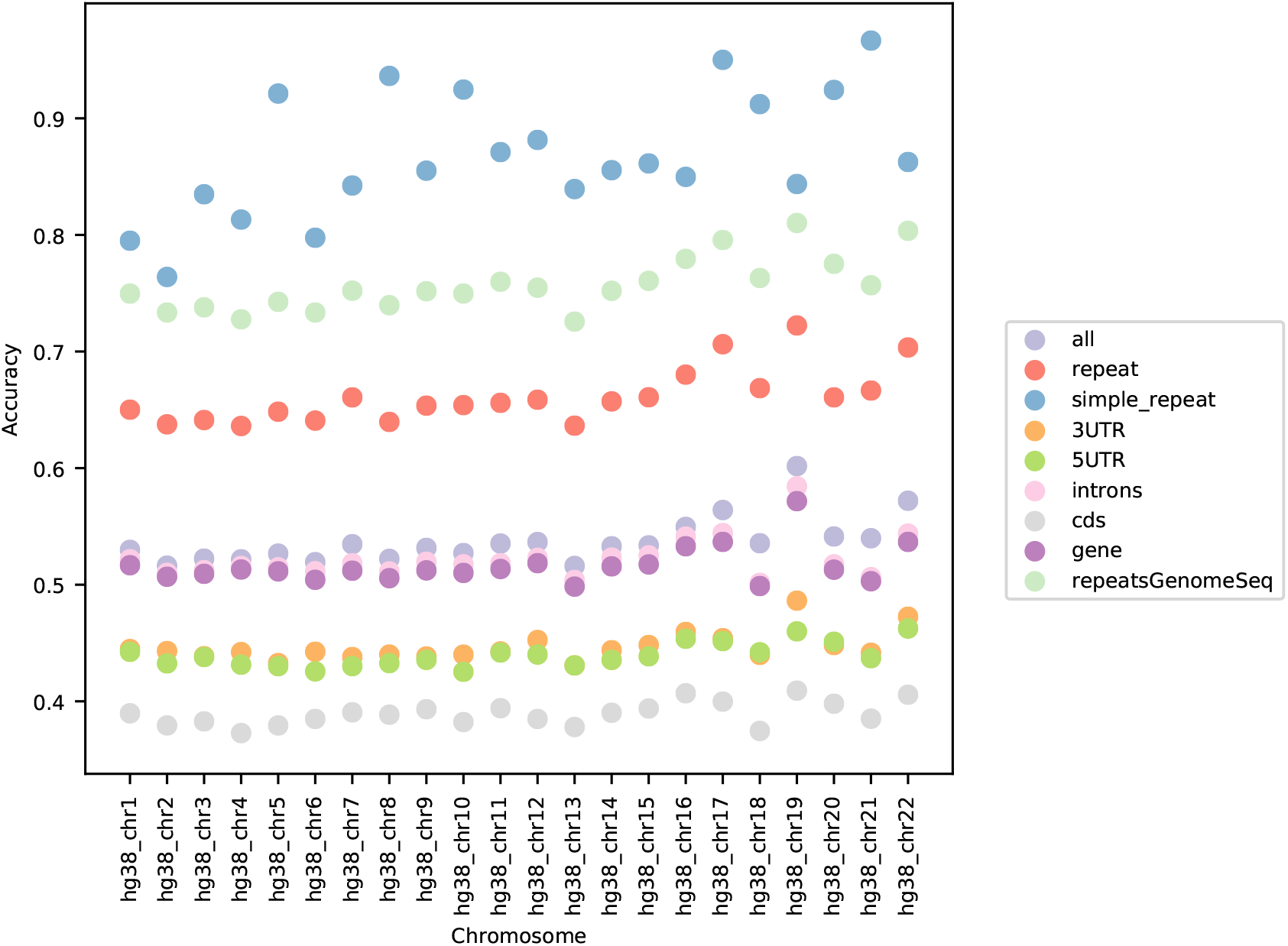
LSTM1 accuracy by annotation per chromosome.

#### Cross-species prediction

Here we use the models to study the similarity of the human and mouse genomes: if the two genomes are highly similar, we expect the performance of a model trained on the mouse genome, when applied to predicting on the human genome, to be very close to that of a human-model. In a general perspective, a measure of genome similarity could be obtained in this way, e.g. by a likelihood ratio test of the models or similarly.

For the concrete task, we trained a model with architecture identical to that of LSTM4 on the mouse reference genome, mm10. We refer to this (trained) model as mouseLSTM4. We return below to describe its performance on the “host” genome (GRCm38/mm10); in this section we consider its performance when applied to the human reference genome, hg38.

As Figure 2 clearly shows, mouseLSTM4 does not perform nearly as well as LSTM4 on hg38. The overall accuracy of mouseLSTM4 lies at about 43%, and varying between about 41% and 48% over the chromosomes (Table 6 in Supplementary Tables and Plots). Remarkably only in the coding parts does mouseLSTM4 perform at par with LSTM4. In other annotated matter the accuracy is lower, with the biggest differences seen in repeat matter having overall accuracy at a mere 46.9%, far from the 63.7% of LSTM4. These values are further supported in Figure 4, which for four annotations shows scatter plots of the probability of the reference-base assigned by the two models, summarized as densities in a grid obtained by binning the two axes. Here two models agreeing would show up as a clear diagonal, while disagreements are seen as bright areas off the diagonal (a pronounced case is seen for repeats, the plot furthest to the right). A bright diagonal and only little disagreement is seen only for the coding sections. Similar plots for chromosome 17, 18 and 19 are placed in the Supplementary Tables and Plots, Figure 1.

**Figure 4:**
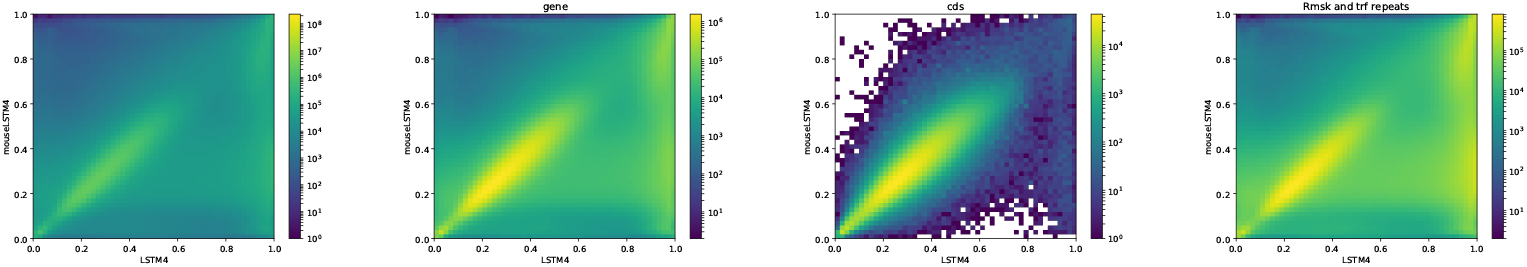
LSTM4 vs mouseLSTM4human. Scatter-plot of probabilities of reference bases in four annotated parts of chromosome 1 of the human reference genome, hg38, according to LSTM4 (x-axis) and mouseLSTM4 (y-axis). From left to right the annotations are: ‘all’ (all positions), ‘gene’, ‘cds’ (coding sequence), ‘repeat masked’.

#### Signals of periodicity in predictions

In addition to examining the models’ performance with respect to the discrete annotations, we looked into whether prediction as a function of genomic position showed signs of periodicity.

In our analysis we segmented each chromosome into adjacent 1 Mb segments and considered at each position the probability assigned by the model to the reference base (the true base); the 1Mb size was chosen for practical reasons. Segments with more than 10% disqualified positions (contexts containing at least one N) and short parts of less than 1Mb were then left out (leaving typically more than 95% of the segments included; the number of segments for each chromosome and other relevant statistics can be found in the Supplementary on Data and data checks). To look for periodic signals we Fourier transformed each of these 1 Mb long arrays of positive numbers (between 0 and 1). In general no frequencies were seen to carry particular weight. However, when taking the norm of all Fourier coefficients within a sliding window of a set amount of frequencies (typically 1000 and a step size of 100 bp) clear patterns emerged as those shown in Figure 5 (similar plots for the other chromosomes can be found in the Supplementary Human Fouriers). To check for the possibility of computational artifacts we shuffled the input arrays and saw no such patterns in the output and similarly when randomizing all positions, while the patterns resisted randomization in half of all positions or in the subset of disqualified positions (Supplementary Methods, Figure 4 and 5). The same patterns found in the adjacent 1 Mb segments also appeared when Fourier transforming randomly picked segments of 1 Mb and other lengths and window sizes (data not included).

**Figure 5:**
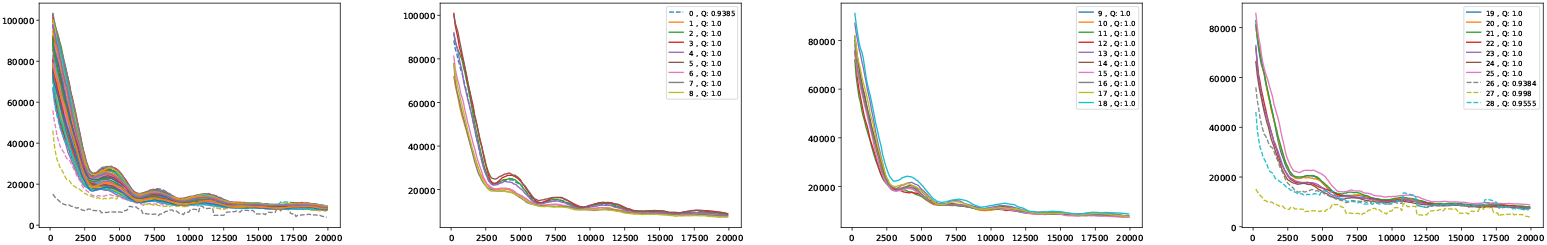
Fourier spectra on the reference-base probability array of LSTM1 for chromosome 20. The chromosome is divided in 64 adjacent segments of 1Mb; segment numbers along with quality fraction are indicated in the legend. First plot (left) shows the results for all segments in this chromosome; the following three plots show the segments covering the first half of the chromosome, including the centromeric region (segments 27 and 28). The plots for all the segments (and all somatic chromosomes) are placed in the Supplementary Fouriers Human.

First, there are peaks at frequencies slightly above 4000, 7500, 11000 and 18000 (and maybe 14000 too). With 1 Mb length these translate into windows of periods of ~ [200 bp, 250 bp], [118 bp, 133 bp], [83 bp, 91 bp] and [52 bp, 56 bp] (and [67 bp, 71 bp]), respectively. If we assign a peak to the mid frequency in a window, the three peaks are at ~ 222, 125, 87 bp and 54 bp (and 69 bp), respectively. These numbers, of which the first and the second are close to the sum of the two following them, appear to be quite close to the lengths of DNA sequences related to nucleosome wrapping: 147 bp wound on each nucleosome and with linker DNA of varying length ~ 20-80 bp ([2] for a recent review).

Second, in all the chromosomes there are segments showing much more abrupt variation. These segments match very well the centromeric regions; in e.g. chromosome 20 (Figure 5) these segments have numbers 27 through 29, which corresponds well with the centromeric region; for more examples see Supplementary Fouriers Human.

These periodicity signals were found not only with the best performing convolutional LSTMs, but also with the *k* = 5 central model and the Markov model (*k* = 14) of [10] albeit with slightly larger span of amplitudes (see Supplementary Fouriers Human, simple models).

In higher frequency ranges we also found signs of periodicity (we only considered LSTM1 here). Thus, in the range 40000 to 140000 (see Supplementary Fouriers Human) a set of three or four peaks occur in all chromosomes, the most conspicuous peak at around 125000 corresponding to a period of about 8 bp.

We wondered whether the signals could have some origin in GC/AT content, i.e. if there are similar signals in GC/AT composition. To that end we applied the Fourier transform to ‘‘boolean arrays indicating Gs and Cs in the DNA”: every 1Mb DNA segment was encoded by leaving a 1 at all G/C-positions and a 0 at all A/T’s. The arrays were segmented and quality filtered as described above (same segmentation). Figure 6 shows an example of the output, clearly suggesting a connection to the signals in the reference-base probability arrays. Peaks are found at similar but somewhat shifted frequencies, and with the weight of the peaks being different from those seen in the models’ prediction arrays (plots of Fouriers on GC/AT content for the other chromosomes can be found in the Supplementary Fouriers Human). Also in the high frequency range 40000 to 140000 this appears to be the case (see Supplementary Fouriers Human); notably the most conspicuous peak in that range is found at slightly less than 100000 corresponding to a period of a little more than 10 bp, close to the ~10.4 period of the double helix.

**Figure 6:**
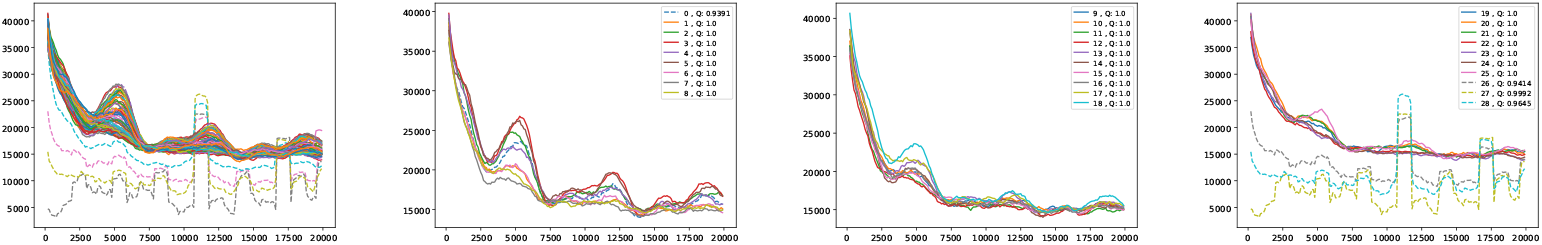
Fourier spectra on the GC-content array for chromosome 20. See caption in Figure 5 for content.

It is also of interest to see if periodic signals occur in the reference-base probability arrays (on hg38) of the model trained on the mouse genome, mouseLSTM4. As seen in Figure 7 this is the case, but with a variation that appears less smooth than in the results from the human model (Figure 5). Comparing further, the two first of these five peaks appear shifted towards lower frequencies, while the following three are placed at similar frequencies; all five peaks have less amplitude, and distributed quite differently than in the human model case.

**Figure 7:**
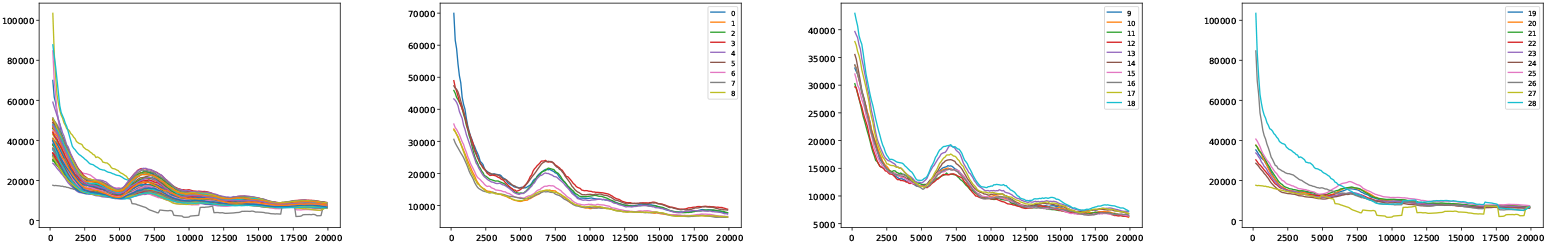
Fourier spectra on the reference-base probability array of mouseLSTM4 for chromosome chr20. See caption in Figure 5 for content.

### Comparison to simpler models

To compare our best neural network model (LSTM1) with the simpler models from our parallel study [10], we worked out likelihood ratio tests [19], all for the null hypothesis of equally performing models. The tests were first done per chromosome by computing the test value on a random sample of 10 % of the positions in the genomic sequence; a plot of the obtained test values can be found in the Supplementary Tables and Plots, Figure 2. A test value for the complete genome was then computed by aggregating the results for the chromosomes. We carried this out for the *k* = 3, 4 and 5 central models and for the Markov model (*k* = 14). In each case the simpler model acted as base (“denominator”) and the LSTM1 model as the “numerator” in the ratio test. The results shown in Table 7 in Supplementary Tables and Plots reveal invariably the superiority of the LSTM1, and with test values increasing with the simplicity of the base model (with p-values essentially all zero a correction for multiple tests is unnecessary). Also, the test values per chromosome increase with the size of the chromosome (and, equivalently, to the sample size) as is evident from Supplementary Tables and Plots, Figure 2, as should be the case on a rejection of the null hypothesis [19].

To illustrate these figures we did model-model plots as in Figure 4, but here for each of the simple models vs. LSMT1 and covering the whole genome (Supplementary Tables and Plots, Fig.3). Recalling that bright off-diagonal areas signify differences in performance, these plots show that LSTM1 generally has higher confidence in its predictions than the simpler models. Thus, for the Markov model, there seems to be a “bend” of the brighter area in favor of LSTM1.

With an overall performance of the Markov model (*k* = 14) of about 52 % [10] it is of further interest to consider how well the model handles the annotated categories as compared to the LSTM1. In Figure 8 we show the Markov model-LSTM1 plots for some major annotations in hg38, chromosome 1 (similar plot for chromosome 17, 18 and 19 can be found in the Supplementary Tables and Plots, Fig.4). The same “bend” of the bright areas is again seen; for the coding sections the bright area even appears shifted above the diagonal, showing that the LSTM1 rather unconditionally is better than the Markov model in the coding regions.

**Figure 8:**
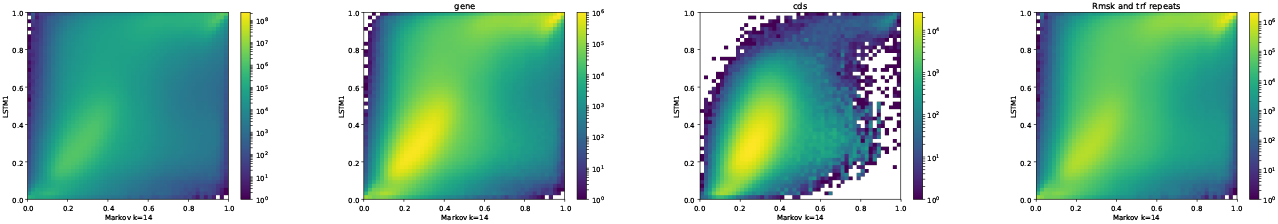
Markov model (*k* = 14) (x-axis) vs LSTM1 (y-axis) on four types of annotated matter in chromosome 1, hg38: ‘all’, ‘gene’, ‘cds’ and ‘repeat masked’ from left to right. Each plot shows the density of the reference-base probabilities according to the named models within the given annotated matter. Please note that colors are not shared between the plots.

In [10] we also applied our models to sets of genetic variants and build a model of mutability. We carried out a subset of the same analyses here, but based on the LSTM1; the results can be found in Supplementary VII.

### Other genomes

We trained and applied models identical to the LSTM4 on reference genomes of yeast (S.cerevisiae), fruit fly (D.melanogaster), mouse (M.musculus) and zebrafish (D.rerio).

#### Accuracy

As reported in Table 1 the obtained accuracy varies much across these organisms. The low accuracy for the yeast-model is quite likely due to the genome size being too small for sufficient training of the model (the number of trainable parameters is too high in comparison). The mediocre performance in the case of the fruit fly can have the very same reason. For the mouse and the zebrafish the performance is quite in line with that for the human genome (LSTM4); the higher level for the zebrafish is apparently partially stemming from repeat sequence (repeats consume about 50 % of the zebrafish genome).

While we did not consider the more narrowly annotated matter, we looked into whether or not periodic signals were found in the predictions also for these genomes.

#### Signals of periodicity: in mouse? in zebrafish?

Broadly, for yeast, fruit fly and zebrafish we found no persistent signals across the chromosomes, either in the reference-base probabilities or GC/AT content. One exception is though a rather clear peak (less in yeast than in fruit fly and zebrafish) corresponding to a period of about 10 bp, i.e. close to the turn of a DNA helix. Most notable apart from this is maybe that in fruit fly and yeast the decay of the coefficients varies substantially across the segments (between two extremes, roughly). More surprisingly, we found no clear persistent signals in the mouse. However, there is some “bundling” at particular frequencies: for reference-base probabilities at frequencies around 5000 (but very weakly) and for GC/AT content (more strongly, but not strong) around 7500 and, maybe, 12500, the segments appear more tightly bundled than elsewhere. Figure 9 shows the plots resulting from the very same Fourier analysis that we carried out for human, but for chromosome 11 of the mouse reference genome, mm10 (results for other chromosomes can be found in the Supplementary section “Fouriers Mouse”). In the higher frequency range (40000 to 140000) there seems also to be some structure in GC/AT content, the same three peaky regions reappearing across the chromosomes as in human (at around 60000, 100000 and 120000).

**Figure 9:**
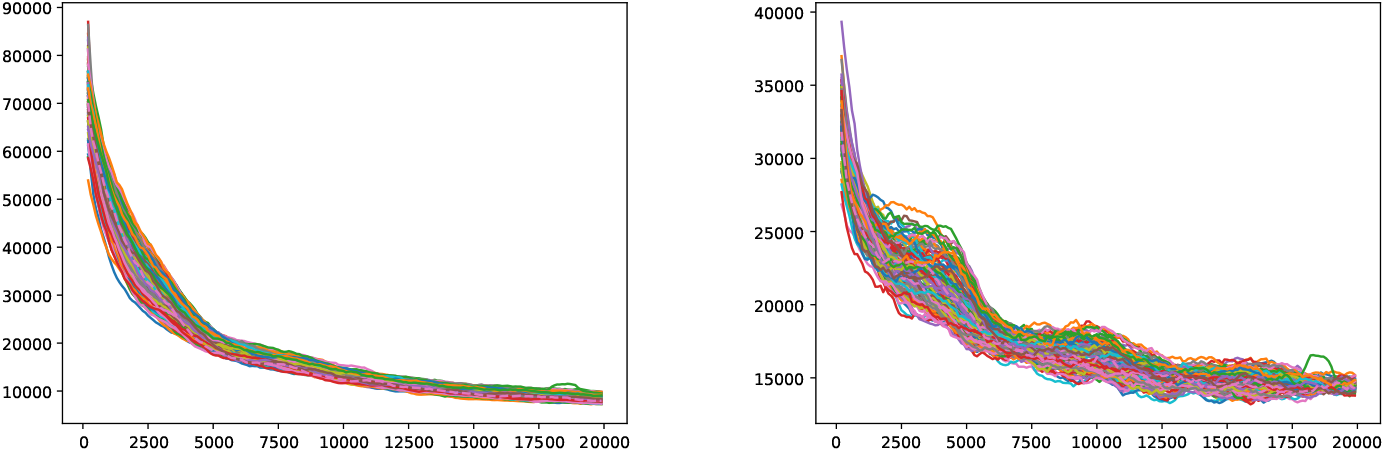
mouseLSTM4 and GC/AT content on mouse genome (mm10). Fourier spectra on reference-base probability and GC/AT content for chromosome chr11. The genome is divided in adjacent segments of 1Mb. Left-hand plot shows the results for the mouseLSTM4 predictions in all segments in this chromosome, the right-hand plot shows the similar results for the GC/AT content.

## Discussion

Our motivation for using neural networks for the task of “predicting DNA from context” was that larger flanks than those within reach by simple models as in [1] [10] could contain information that would allow improvement of the prediction. For each *k* = 3, 4, 5 the central model [10] provides an upper bound for the overall accuracy using flanks of size *k* and our neural networks outperform these clearly. Also, LSTM1, which uses flanks of size 200, outperforms the flanks-50 LSTM4 (these models are almost identical). So our main hypothesis seems right. To further support this we earlier ran tests shuffling and randomizing parts of the contexts, and saw the expected decrease in performance (data not included). However, while our models break the 50% mark, it is seemingly not by much, and our very best neural net adds only about 2% points to the accuracy of the Markov model (*k* = 14). We have though noted that obtaining these extra few percentage points appears to be a struggle; about two thirds of the complete training of LSTM1 are spend there. These extra percentage points are obtained across the structural parts (annotations), and not only by improving the performance e.g. in repetitive sequences (compare LSTM11 and LSTM1 in Figure 2). Further, the likelihood ratio tests show that the LSTMs are superior to the simpler models, with clear significance. In terms of number of parameters, one facet of model complexity, the LSTM1/4 is the lightest with about 2.3 million; the *k* = 5 central model has about 3.1 million parameters and the Markov *k* =14 model about 0.8 billion. If the aim is not to improve model performance, the Markov model comes out as the model of choice, due to its simplicity, good performance and ease of use. As opposed to the neural networks, the performance of the Markov model is though ‘saturated”. And our experiments do of course not show that our “hi-score” (about 54 %) is a bound that neural nets cannot push; to the contrary it would be surprising if that would not be doable.

Considering the level of the accuracy obtained on the human genome, it does catch the eye that it ends up close to 50%, and we have noticed that this level is close to the accuracy reached on the “gene” and “intron” annotated part. This probably reflects that introns dominate these regions and their composition is very similar to intergenic regions. Looking into how the accuracy was obtained, we expected that the prediction would be easier in repetitive sequences than elsewhere. This is clearly the case, but in all (other) annotated parts the performance is far better than by random guessing (Figure 2). Variation across chromosomes (Figure 1) and chromosomes/annotations were rather small (Figure 3).

When training and running models on other (reference) genomes we saw performance at the same level as that for the human genome (mouse, zebrafish), while for smaller genomes lower accuracy was had (fruit fly, yeast). We did not try to explore the sources of the performance for these genomes as far as we did for the human genome. Repetitive sequences though plays a similar role. For all these cases we used the architecture of the LSTM4; thus the trained models were identical except for the values of the trainable parameters. This allows direct comparison, but the amount of training had to be kept much lower for the small genomes (fruit fly, yeast) than for the larger genomes (human, mouse, zebrafish), which could all be trained to similar extent. The modest performance in the cases of yeast and fruit fly, even when allowing the training to cover the genomes several times, suggests that the number of data points is too low as compared to the number of parameters. This calls for experiments using more economic models in terms of size/number of parameters.

Having models trained on different organisms opens a possibility of comparison of their genomes. We found it particularly interesting to apply the model trained on the mouse genome (mouse-model, mouseLSTM4) to the human genome and then compare to the human-model (LSTM4). This gave the rather surprising result that apart from the coding regions, the mouse-model was performing substantially worse than for the identical model trained on the human genome (LSTM4). The congruence on the coding part hangs well together with the known genetic similarities between mouse and human. The differences can have many sources. For one, the repetitive sequences in the two genomes may be very different. And certainly, as the models fall long short of predicting their host DNA perfectly, their “representation” of that DNA may have large imperfections, and possibly specific to the DNA in question. At the extreme, comparing two essentially random models would be illogical; however, our models are clearly very far from such a state.

That this is indeed the case is well supported by the obtained accuracy and the likelihood-ratio tests that we have carried out, but also by the periodic signals that we have found. In the case of the human genome, but hardly in mouse, the periodicities in the reference-base probabilities seem connected to those seen in the GC/AT content. In the human case the periods in these are very similar and in the mouse the same kind of “bundling” appears in both (though at somewhat different frequency ranges, and only very weakly in the reference-base probabilities). This reveals that, at least to some extent, the model is capable of capturing this structure. The differences between the GC/AT content and the models’ predictions can have several sources: the model has not picked up perfectly the GC/AT composition bias; there may exist other periodic patterns to which the model has adapted.

Regarding the method, applying Fourier transformation to DNA sequences is certainly not new. Thus e.g. in [18] and maybe earlier, Fourier transform was applied to DNA sequence. Our approach can be summarized as considering “the cumulative power spectrum in a running window”. A very recent application of the Fourier transform to DNA and aimed at genome comparison, consists in considering the cumulative power spectrum of each of the indicator functions of the four bases [5] [13]. In between [18] and these publications much has though been done in spectral analysis of DNA sequences by Fourier transforms, see e.g. [11] [20] and references therein. The well-known ~10.4 bases periodicity in dinucleotide composition which has been linked to histone binding, has also called for Fourier analyses but aimed at di- or trinucleotide composition [15] [20] [21]. The work [11] includes also a rigorous statistical analysis for significance in spectra, something we have not touched upon. Our findings are obtained by visual inspection. As mentioned, analyses similar to those of GC/AT content that we have carried out and reported here we have not come across in the literature. To rule out the possibility of mere computational artifacts, we did simple shuffling and randomization tests. Also, the periodicity signals vary substantially over the organisms considered. The patterns seen in GC/AT content rule out a modeling artifact, as does the fact that the reference-base probabilities of the Markov model show almost identical signs of periodicity as for the LSTM1 on human (chromosome 22). Finally, in human the characteristic patterns over the segments were seen to change abruptly in centromeric regions; in mouse this phenomenon appeared absent well in line with the fact that centromeres in mouse are found at the ends of the chromosomes with essentially no short chromosome arm.

We aim to dedicate a paper to exploring a possible biological explanation for these patterns, but some treatment is called for here. The periods that we have found in the case of the human reference genome could point to the chromatin’s “beads-on-a-string” structure, in which the DNA is wound up over nucleosomes at quite regular spacing. This structure is though probably local, dynamic, and may vary much across a chromosome. Notwithstanding, we found clear signs of periodic regularity, with periods shared by all autosomal chromosomes. The similarity to the periodic signs in GC/AT content could support this [8], as do the abrupt changes in the patterns in segments covering centromeres (human) [14]; see also the more recent review [4].

On the other hand, patterns with these periods showed up only in the human genome. In mouse there is a certain “bundling” in the GC/AT content case, but only very vaguely in the reference-base probabilities. For the other organisms’ genomes that we considered, signals were essentially absent, except for a common, though not equally clear, peaky region corresponding to periods of about 10 bp. These hint at the well-known ~ 10.4 bases periodicity in dinucleotide occurence, accepted to be connected to nucleosome positioning. But the clear, longer periods only appear in human, which seems to undermine our nucleosome-positioning hypothesis, since nucleosome organization is regarded to be quite similar across species. These periodicities are hardly multiples of the ~ 10.4 bases repeat pattern: Were this the case, a series of corresponding frequencies should be expected, which appears quite clearly not to be the case. Also, in the other genomes we should then have observed such ‘harmonic series’. So possibly the signals in human have other sources or an in-between — mechanisms of nucleosome positioning may differ across species.

Finally, we can, strictly speaking, only say that we have seen these signals in a reference genome. But a complete lack of a biological source seems highly unrealistic. At the other extreme, if the periodic patterns in the human genome are connected to nucleosome positioning, it could be a piece in the puzzle of the gene regulation in non-coding material in humans (and possibly for other species). More broadly, since these patterns appear to be different between species, they could very well be of benefit in several areas.

## Methods and implementation

See the dedicated Supplementaries. We intend to make the code used for this paper available on github.

### Data

For all training, testing and prediction we used reference genomes downloaded from major resources [12] [16]. For more details on the downloads along with some statistics and checks of the data see Supplementary Data and data checks.

## Supporting information

Supplementary: Data and datachecks

Supplementary IIa

Supplementary IIb

Supplementary IIc

Supplementary IId

Supplementary IIe

Supplementary VII

## Acknowledgments

This work could not have been done without access over a considerable period of time to servers with ample storage space and fine GPU facilities. It is therefore a great pleasure to have the opportunity to thank Robin Andersson and Albin Sandelin at the Department of Biology of the University of Copenhagen for generously allowing us to run our analyses on their GPUs virtually *ad libitum* and to Hanne Munkholm for server hospitality and support to make it possible in the first place. CG also wants to thank Nicolas Alcaraz for help with the GPU set up. It is also a pleasure to thank Piero Fariselli for initial discussions and for sharing his material with us.

## Author contributions

AK and CG conceived of the study. CG designed the neural networks with assistance from AK and DE. CG wrote and ran all code, except a speed up by DE. YL did data management. CG drafted the paper. All authors revised and approved the final text.

## Supporting Material

The following supplementary materials (pdf files) are available:

1. Supplementary: Methods
2. Supplementary: Data and data checks
3. Supplementary I: Tables and plots
4. Supplementary II, a, b, c, d: Fouriers, human genome; LSTM1 (a,b), GC/AT content (c,d)
5. Supplementary IIe: Fouriers, human genome, simple models (chr22); mouseLSTM4 (chr20)
6. Supplementary III, a, b, c, d: Fouriers, mouse genome; mouseLSTM4 (a,b), GC/AT content (c,d)
7. Supplementary IV: Fouriers, zebrafish genome; LSTM4 (a,b), GC/AT content (c,d)
8. Supplementary V, a, b: Fouriers, fruit fly genome; LSTM4 (a), GC/AT content (b)
9. Supplementary VI, a, b: Fouriers, yeast genome; LSTM4 (a), GC/AT content (b)
10. Supplementary VII: SNP analysis

