## Supplementary: Methods for "Prediction of DNA from context using neural networks"

April 24, 2021

### Methods

#### Neural networks

##### Training and testing

Neural networks take numeric input and apply arithmetic operations to it to produce a likewise numeric output. To make a neural network applicable to DNA strings, it is necessary to map the four bases (and any "wild cards") to the numeric world. For this we use "one-hot encoding" of the four letters as follows: A is mapped to (1,0,0,0), C to (0,1,0,0), G to (0,0,1,0) and T to (0,0,0,1). Thus the four letters are replaced by four linearly independent vectors in four dimensional real space ($\mathbb{R}^4$); this independence is important for not introducing numeric correlation between the four letters. The encoding is extended to strings of (the four) letters by applying it to each letter in the string. Thus a string of length L is mapped into 4*L dimensional real space ($\mathbb{R}^{4L}$). The computations carried out in the first layer of a network applied to such strings therefore take place in this space.

The input to a model consists in a one-hot encoded genomic context (two flanks, merged or not). The output of our networks consists in a four dimensional real vector following the same encoding and for which the entries are positive and sum to 1. The output can then be interpreted as a probability distribution over the four letters; so for instance the first entry is the probability of having the letter A given the input. This property of the output is had by letting the final layer of the network have a "soft-max" function as its activation. Activation functions are elsewhere set to the "rectified linear", sometimes referred to as ReLu.

To train one of our networks it is fed with pairs of (context, label) where label is the base at the (mid) position of the context. Formally, if we let $x$ denote the genome sequence, for position $i$ and a context of flank size $k$

$$context_i = x_{i-k} \dots x_{i-1} x_{i+1} \dots x_{i+k}$$

and label = $x_i$.

Training sets of such pairs, (context, label), were obtained by sampling from the entirety of the given genome (for convenience e.g. the first 3 billion positions of the human genome); the model named LSTM4S was though only trained on the odd-numbered chromosomes of hg38. The training was run in ”rounds” of 5 million samples; at the end of each round the model was validated at that stage on a similarly sampled set of 1 million contexts. The full training consisted in completing 200 such rounds (for the smaller genomes of yeast and fruit fly we used fewer rounds, see more later). By the sizes of the genomes (human, mouse, zebrafish) of about 2.5-3 billion positions and the size of the complete 200 training rounds of about 1 billion positions (each round being 5 million), the probability of a given position being included in the training was about one-third to two-fifths. Thus it seemed hardly necessary to keep validation and training sets disjoint (the former are used for the validation of the model after each training round; the real test of a model is had when applying it to new data, which here consists in about two-thirds to three-fifths of an entire genome). However, for all models except the LSTM1, the training and validation sets were separated by a 80-20 split: at initiation of the training 80 % of the total genomic positions considered were picked randomly as sampling material for the training sets, and the remaining 20 % were left for sampling the validation sets. That the validation during training of the LSTM1 is just as reliable as for the other models is clear from Figure 1, which also reveals that no overfiting was present. Indeed, the final tests — applying the trained models to the full genomes — showed accuracies close to those obtained in the validations. To clarify that the final prediction on the full genomes gives a fair estimate of the performance (not exaggerated due to training material being included in the test material or, more generally, overlapping context in the training and test sets), we trained a model, LSTM4S, only on the odd-numbered chromosomes of hg38. The architecture of LSTM4S ’lies in between’ that of LSTM4 and that of LSTM1 (more below). The performance of this model turned out to be only marginally better than that of LSTM4 (Fig. 2 and Fig.3 below), and with no signs that the generalization to the even-numbered chromosomes is out of level with the performance on the odd-numbered chromosomes (results of LSTM4S are included with those of the other models in tables in the Suppl. tables and plots).

At some genomic positions the true base is unknown or uncertain, and a "wild card", typically the letter N, occurs at such positions. If a context or its label contained a wild card it was disqualified from the training.

The training of the LSTM1 was special in one further aspect. The trained model ought to have the following "symmetry" property: when applied to the reverse complement of a given context, its output must be the (reverse) complement of the original output. Formally, if $model(c) = b$ where $c$ is a context and $b$ is the output base (label), then

$$model(\bar{c}) = \bar{b}$$

where $^{-}$ means reverse complement. To guide the training of a model towards having this property, one can let the training batches be "invariant under reverse complement": each round of 5 million samples was in fact obtained in 100 "epochs" each containing 100 steps of optimizing the model's parameters; each step in turn comprised a batch of 500 sampled contexts with corresponding labels. By augmenting each batch by the reverse complemented contexts and labels, the parameters were continuously updated on batches of 1000 samples, each invariant under taking reverse complement. For LSTM1 each training round then in fact counted 10 million samples, and the full training session ran therefore through 100 rounds and not 200.

For each full training session the model showing the best validation result during all rounds were picked out and used for further analysis. In all cases the best round was found in the late stages of the training (i.e. close to the final round). The only exception to this was LSTM11, which is simply the model obtained at repeat (round) 15 of the training of LSTM1.

For the small genomes (yeast and fruit fly) we let the training continue far beyond only covering a relatively small fraction of the entire genomes, and then also considered models obtained early in this training as well as models at much later stages. For both species we split the input genome sequence in a training and a validation pool (80-20 $\%$) as described above. Thus for yeast we considered a model, called LSTM41, obtained after the first repeat (5 million samples, slightly less than half the genome size) and a model, named LSTM4, obtained at 59 rounds/repeats (where the process will have covered the 80 $\%$ of the genome designated for training about 30 times). For the fruit fly we considered an early stage model, LSTM41, at 12 repeats (the training material then covering about one third of the genome) and a late stage model, LSTM4, at 168 rounds/repeats (where the process will have covered the training material about 6 times). The results we report are for the late stage models unless otherwise stated (the results for the early state models can be found in the Suppl. tables and plots). The results for the late stage models cannot be seen as predictions; the models represent our ”best fits” to the data and the reported accuracy is a measure of the quality of the goodness of the fit. (The aim of the paper is not really to predict the DNA, but rather to fit models with the purpose of investigating – or revealing — structure in the DNA strings. The aim of prediction serves as a good frame in which to do this. But we could have trained the models on e.g. the human genome far more extensively, while monitoring over-fitting).

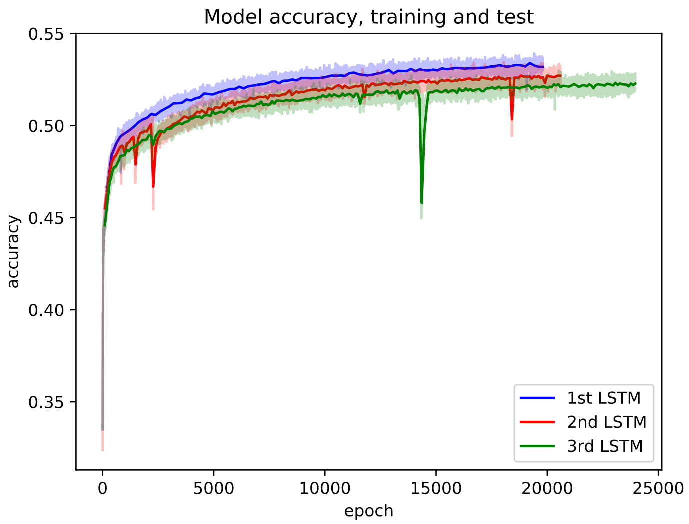


Figure 1: Accuracy during training (dimmed) and validating (solid line) of the best convolutional-LSTMs (1st LSTM is the model referred to as LSTM1 elsewhere in this text; the two other models were similar to LSTM1).

#### Architectures

We experimented with various types of architectures and settings, training all on human DNA (assembly hg38).

First we tried classic feed forward networks (best accuracy of about 46 %) and then convolutional networks (best accuracy of about 48 %). We also experimented with merging a network and a central model, resulting though in unstable training (data not included).

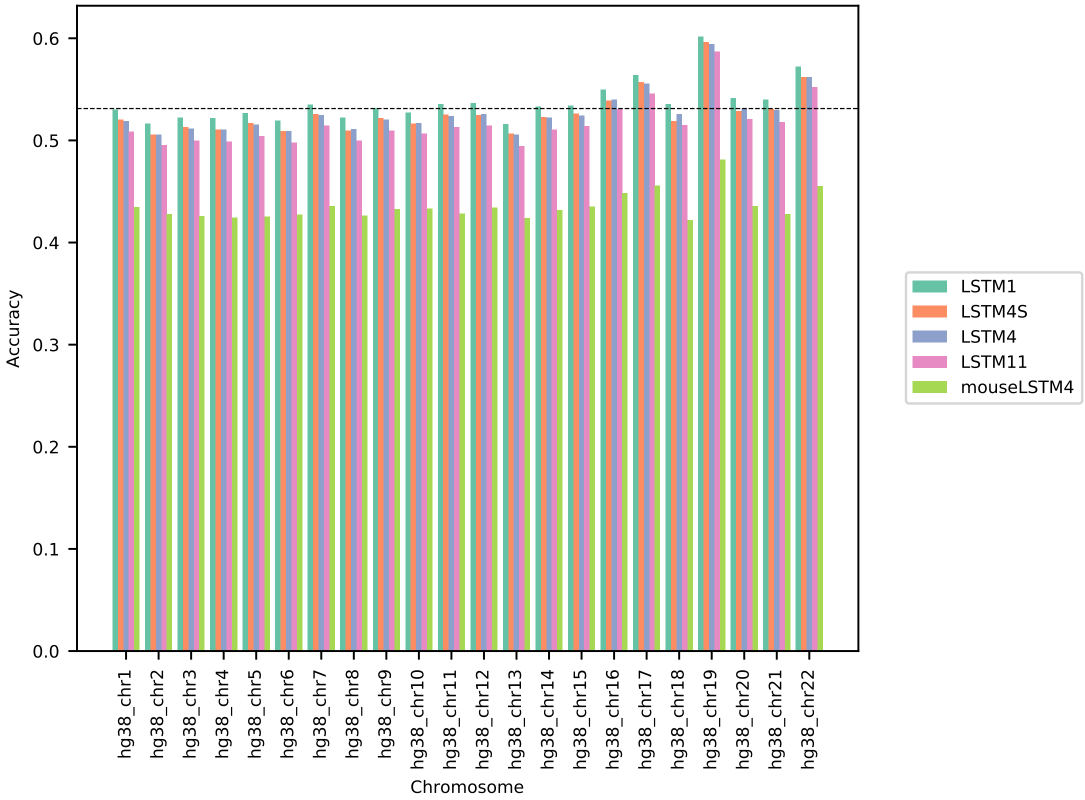


Figure 2: Model accuracy comparison across the human autosomal chromosomes (hg38). The named models are described in the main text (for LSTM4S see also above). The dotted line indicates the average performance of LSTM1. All values can be found in Table 1 in the Supplementary Tables and Plots.

Upon moving to Long Short term Memory (LSTM) networks [1] accuracy could be improved to pass the 50% mark. In all these the LSTM layers are applied bi-directionally: moving forward (5' to 3') on the left-hand flank part and backward on the right-hand flank part. With the aim to include a kind of "word encoding" we though let the networks be initiated by one or two convolutional layers, with filters of size 3 or 4 and applied with a stride of 1. The number of filters were very simply inspired by the number of words in the four letter alphabet of length equal to the filter size. Following this word encoding, which is applied separately to the two flanks, the LSTM layers follow (mostly just two layers).

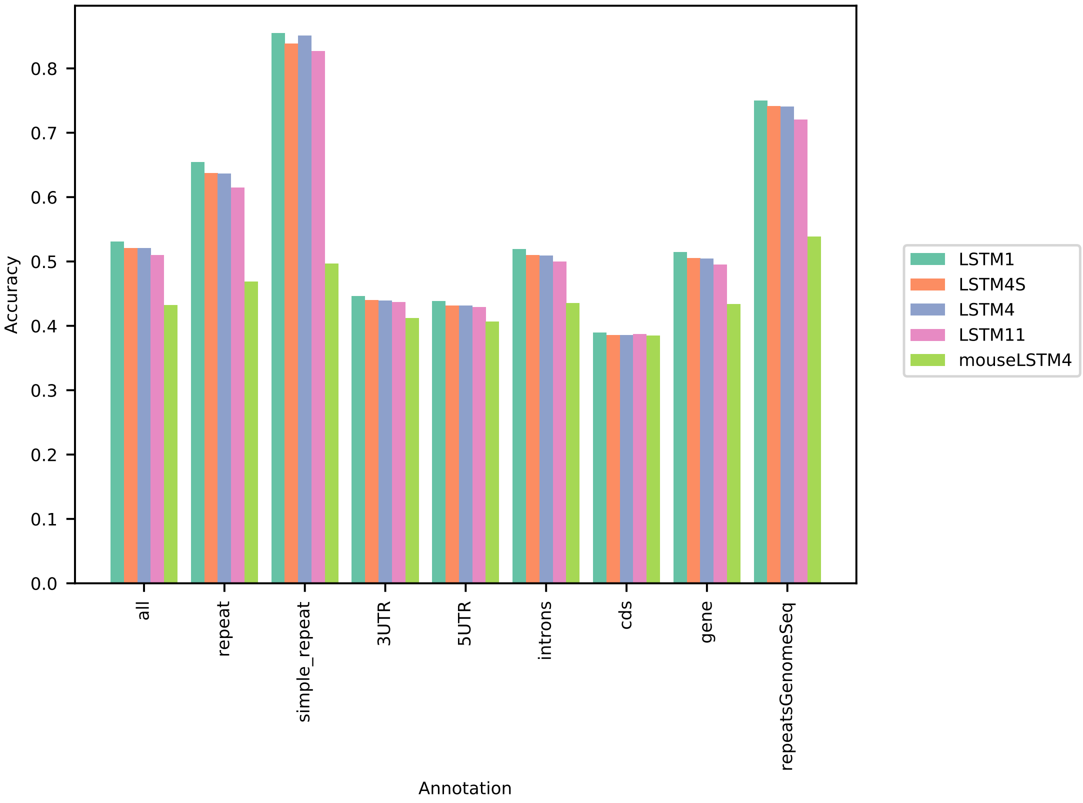


Figure 3: Five model accuracy comparison by annotation. All values can be found in Table 2 in the Supplementary Tables and Plots.

The final LSTM layer outputs its final node value from the left-hand flank part and from the right-hand flank part. These are concatenated and fed into a final dense layer, intended for the purpose of obtaining a condensed representation of DNA context. The final layer outputs four positive real numbers summing to 1 as described above.

The architecture of the model that we refer to as LSTM4 is laid out graphically in Figure 4 . The architecture of the model referred to as LSTM1 is almost identical to this model, except that it has an extra word encoding convolutional layer and uses flanks of sizes 200 to either side, shown in Figure 5 . The model called LSTM11 is identical to LSTM1; LSTM11 is simply the model taken out after 15 repeats of the training for LSTM1 (similarly the yeast and drosophila LSTM4 models LSTM41 are obtained early in the training of the LSTM4's). The model named LSTM4S has the same convolutional layers as LSTM1, but uses contexts of flank size 50 (as the LSTM4), so is an LSTM4-LSTM1 intermediate.

As for the number of trainable parameters these three models have about 2.3 million.

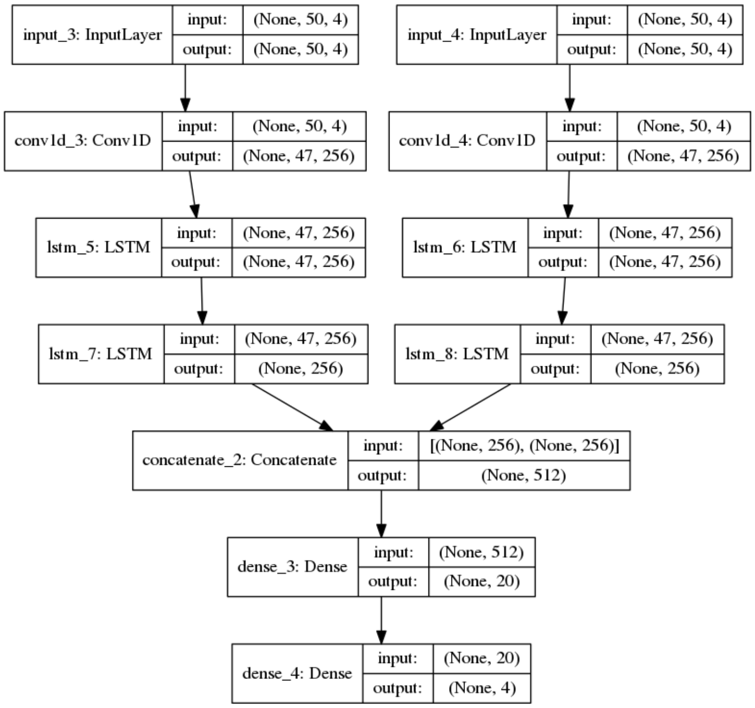


Figure 4: Our convolutional-LSTM LSTM4. Flanks have size 50 and the LSTM layers are applied bi-directionally on the flanks, going left-to-right on the left-hand flank and vice verse on the right. The first LSTM layers each output a sequence of the same length as the input; the second LSTM layers each output the final position.

#### Implementation

We implemented the neural networks using Keras/Tensorflow [3], an open-source machine learning platform in Python. Among many great facilities, this includes automatic off-loading of core computations to Graphical Processing Units (GPUs), making the training process and predictions doable on a timescale of days, weeks

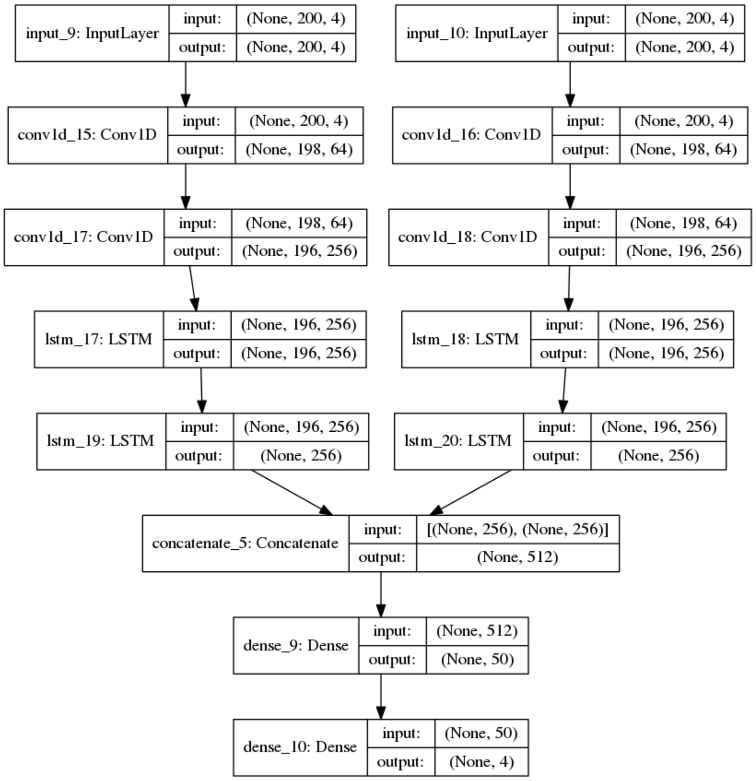


Figure 5: Our convolutional-LSTM LSTM1. Similar to LSTM4 (above), but flanks have size 200 and the word encoding consists of two convolutinoal layers, the first encoding three letter words.

or months. Our server ran with 16 GB RAM dedicated to the GPU, a Nvidia Tesla P100.

The training time varied considerably: for the simpler models (feed forward and convolutional) the training on the human genome (as described above) took one to two days. The convolutional-LSTMSs took much more time to train, but severely influenced by the flanks size: for flanks of size 50 (LSTM4) the training took about 1 week; for flanks about 200 (LSTM1) some 3 weeks were spend. We observed that the last 2 weeks of training resulted in a gain in accuracy of about two percentages points.

##### Prediction

After training, the model was applied to a complete genome (usually the one on which it was trained): for every position the model was fed with the context and the four probabilities output by the model was kept (this was carried out in batches of 528 positions). The base predicted by the model was then set to the letter having the highest probability: at genomic position $i$ the predicted base in one-hot encoding is

$$prediction_i = \mathbb{1}_{argmax_{j=1,2,3,4}(p_i(j))}$$

where $\mathbb{1}$ is short-hand for a "boolean array on four entries" (so e.g. $\mathbb{1}_3$ is $(0,0,1,0)$ i.e. the encoding of letter G).

For disqualified positions (i.e. if the context or label contained a non-ACGT letter — a wild card) the prediction was set to a (different) wild card. Disqualified positions were recorded and kept out of downstream computations, e.g. of accuracy and likelihood ratio tests.

A model's output (array of predictions) consists in an array of length equal to the covered genome string. For handling this, we partitioned the input genome string (from which a context is build at each position and input to the model) in adjacent, disjoint 1 Mb segments, as many as could fit into the genome string. Thus all output (predictions) and information related to it (e.g. a boolean array recording if a position was qualified or not) were segmented in the very same way.

For a few of the human chromosomes a large heading section was disregarded, since it consists in N's (e.g. for chr22: the start position was set to 10500000; for more see the Supplementary on Data and data checks).

##### Fourier transformations

Fourier transformation was done on every 1 Mb segment for which more than 90 $\%$ of the positions were qualified. The transformation was carried out on the model's output and on arrays encoding GC/AT content. In the first case, the input to the transformation was the model's probability of the reference base at every position (the reference base being the base in the genome at a given position). For the GC/AT content case, the input consisted in arrays having a 1 at every G and C, and zero elsewhere (also in 1 Mb segments). These arrays had then no bearing on the models, but followed the same segmentation.

Since the input arrays have only real-value entries, the Fourier transforms are Hermitian: the coefficient of a given frequency is the complex conjugate of the coefficient for the corresponding negative frequency. For this reason we only show the (norms of) the coefficients in the positive frequency range.

In general the Fourier coefficients turned out not to reveal any structure that met the eye (an example is shown in Figure 6, left). However, when summing up the 2-norm of the coefficients within a sliding window (in "frequency space" that is), peaked patterns occurred in several cases (an example is shown in Figure 6, mid). This amounts to taking the 2-norm of the part of the input function that gives rise to the Fourier coefficients at the frequencies within each window (apart from a factor of the square root of two: for a given window we ought to include also the corresponding window of negative frequencies, but the 2-norm of these two parts equals by Pythagoras' theorem and the Hermitian property of the transform the square root of two times the 2-norm of the coefficients in one of the two windows). A window size of 1000 and a step size of 100 were generally used (larger genomes), while shorter for yeast (window size 100 and step size 10).

The plots of the Fouriers transformation results are truncated for the very first low frequencies and from high frequencies; thus we show plots covering various frequency ranges, e.g. [200, 45000] and [40000, 120000]. The reason is that the left out parts appear uninteresting and if including them the signals would "drown" somewhat in the scale.

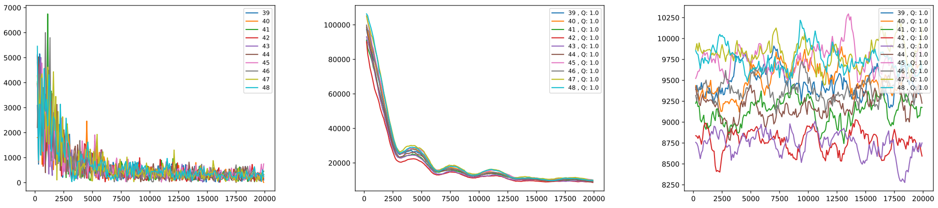


Figure 6: Plots of outputs from Fouriers transforming the reference-base probability for segments 40 to 49 (1 Mb each) of chromosome hg38_chr19 according to LSTM1. From the left: absolute values of the Fourier coeffcients; L2-norm of Fourier coeffecients in a sliding window of 1000 (freqeuncies); absolute value of the Fourier coeffecients had when shuffling the input. The segment numbers along with the fraction of qualified positions are indicated in the legend.

To check that the obtained patterns were not numeric artifacts, we randomized the input arrays in several ways. First, input arrays were shuffled; upon Fourier transformation no patterns were seen (Figure 6, right). Second, we whitened (random values in $[0, 1]$) various parts of the arrays: all disqualified positions; 50

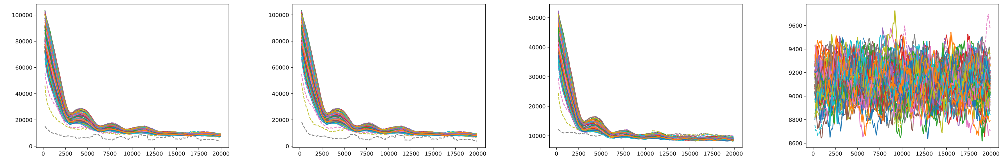

Figure 7: Plots of outputs from Fouriers transforming the reference-base probability of chromosome hg38_chr20 according to LSTM1, as well as whitened as described in th text. Furthest left, the untouched L2-norm of Fourier coeffecients in a sliding window of 1000 (freqeuncies). Then follows the whitened versions: randomized at all disqualified positions; next at 50 % randomly picked positions and finally at all positions. The segment numbers along with the fraction of qualified positions are indicated in the legend.

% randomly picked and 100 % of all positions (Figure 7). Randomizing at all disqualified positions has only very little effect (which of course only hits the segments with some disqualified positions). Randomizing all positions wiped out the signal, while the signal survived whitening even as much as half of all positions, albeit weakened. We made similar observations for Fourier output on the GC/AT content. This clearly points to the period signals are not computational artifacts. The differences seen in the results found for the genomes of various organisms also underpin this. Further, dramatic differences in patterns were seen in some few segments of chromosomes of the human genome (hg38); these segments appeared to overlap quite precisely with centromeric regions. This behaviour was not seen in the case of the mouse genome, fitting well with the fact that in mouse the centromeres are placed at the ends of the chromosomes (the short arms being essentially absent).

### ”Simple” models

We tested our neural networks (the LSTMs) against the ”simple models” of [2]. For ease of implementation we ran the $k$ central models for $k = 3, 4, 5$ to serve this paper (the $k = 5$ central model interpolated from the $k = 4$ central model); the results could easily be stored in Python dictionaries. The run of the Markov model $k = 14$ was though taken from [2], the results being read into Python from a flat file.

#### Likelihood ratio tests

Likelihood ratio tests for non-nested models [4] were carried out for the LSTMs vs each of the simpler models. We also implemented a nested-models version. The tests were implemented in Python; we checked our implementation by comparing the non-nested model test to a nested-model test, when both were applied to the nested-models case (two $k$ central models).

The likelihood ratio tests of LSTM1 vs. each of the simpler models were carried out on each chromosome of the human genome GRCh38; each of these tests ran on a random sample of 10 $\%$ of the chromosome's positions. Results were recorded so that a test figure for the entirety of the chromosomes could be computed too. This allowed also to see if the test figures' behaviour with respect to sample size was as expected [4] (upon a rejection of the null-hypothesis of equally well performing models, the test size should go to $+\infty$ for sample sizes going to $+\infty$ in the case that the "numerator" model was superior). For a plot of the test sizes per chromosome see the Supplementary Plots and Tables.
