## Supplementary: Data and datachecks for "Prediction of DNA from context using neural networks"

November 26, 2020

### Genome data

Reference genomes for human, mouse, fruit fly, zebrafish and yeast were downloaded at public sites. Below we give details for each organism along with some statistics useful in conjunction with the predictions and accuracy computations for which we have reported results.

#### Human

We downloaded assembly GRCh38 from [2] in ”soft masked” format at the site https://www.ncbi.nlm.nih.gov/genome/?term=human (the same sequence can be found as hg38 on [3]). Only the primary sequences were used. Sequences for the individual chromosomes (chr1-chr22) were obtained by splitting the full sequences. The (soft) repeat masking is done by the WindowMasker (the hg38 sequence at [3] probably has a slightly different repeat masking). We refer to this repeat matter as ”repeat masked” or ”RepeatGenomeSeq”.

The files (bed-format) we used for annotations (simple repeats, repeats, UTR’s, gene and coding section, cds) were downloaded from [3] at the interactive site https://genome.ucsc.edu/cgi-bin/hgTables.

To check the hg38 chromosome sequences we got in this way (and which we have used in all our analyses) we compared them base-by-base to the ones that can be had directly from the UCSC genome browser [3]. Concretely we downloaded the sequences of the individual chromosomes that we used for checking at http://hgdownload.soe.ucsc.edu/goldenPath/hg38/chromosomes/ (July, 2020).

In a second test aimed to check the soundness of our encoding, we report the number of different bases found upon inverting the one-hot encoded chromosome strings back to the genomic alphabet.

The results of these two tests can be seen in Table 1, which reports the differences in number of bases. Discrepancies are seen but so few that they cannot harm our analyses.

Table 2 shows the ”statistics” of the input to our predictions and accuracy computations. Qualified positions are those for which a context given by 50 bases (LSTM4 setting; for LSTM1 we used 200 bases) to both sides plus the position itself does not contain a non-ACGTacgt letter. The ”fraction of qualified” is the fraction of qualified positions relative to the whole genome sequences (for each chromosome). The number of segments is the number of segments (here 1 Mb long) for which predictions were made. For a few chromosomes (13, 14, 15 and 22) the number of segments does not match the length of the genome sequence; for these chromosomes there are long initial stretches of N's and we chose to not do predictions on them (all these positions are disqualified so the prediction is irrelevant). Therefore, for these chromosomes, the predictions start at positions later than 0; the start positions were chr13: 16 million, chr14: 16 million, chr15: 17 million, chr22: 10.5 million. The low number of segments for these chromosomes reflects covering only these ”truncated” sequences. These four chromosomes also show the lowest fraction of qualified positions, which though overall is high (generally above 90 %).

The remaining numbers in Table 2 give the fraction of the qualified positions having the named annotation. These were used for computing the accuracy of the prediction restricted to these annotated parts of the genome.

Finally, for the human genome, Figure 1 and Table 3 show the GC content per annotation type. The coding sections (cds) and 5UTRs have the highest GC content; in the whole genomic sequence the fraction is about 41 %.

### Mouse

The soft-masked mouse genome assembly GRCm38 (mm10) was downloaded from [1] at site ftp://ftp.ensembl.org/pub/release-101/fasta/mus_musculus/dna/. As with the human genome the file was split into files (sequences) for individual chromosomes.

| chr | length | #diffs 1-hot | #diffs bases |
|---|---|---|---|
| hg38_chr1 | 248956422 | 2 | 2 |
| hg38_chr2 | 242193529 | 7 | 9 |
| hg38_chr3 | 198295559 | 5 | 7 |
| hg38_chr4 | 190214555 | 0 | 0 |
| hg38_chr5 | 181538259 | 0 | 0 |
| hg38_chr6 | 170805979 | 1 | 1 |
| hg38_chr7 | 159345973 | 4 | 4 |
| hg38_chr8 | 145138636 | 0 | 0 |
| hg38_chr9 | 138394717 | 3 | 3 |
| hg38_chr10 | 133797422 | 30 | 36 |
| hg38_chr11 | 135086622 | 0 | 0 |
| hg38_chr12 | 133275309 | 3 | 3 |
| hg38_chr13 | 114364328 | 3 | 3 |
| hg38_chr14 | 107043718 | 0 | 0 |
| hg38_chr15 | 101991189 | 0 | 0 |
| hg38_chr16 | 90338345 | 1 | 1 |
| hg38_chr17 | 83257441 | 11 | 12 |
| hg38_chr18 | 80373285 | 0 | 0 |
| hg38_chr19 | 58617616 | 0 | 0 |
| hg38_chr20 | 64444167 | 0 | 0 |
| hg38_chr21 | 46709983 | 3 | 3 |
| hg38_chr22 | 50818468 | 4 | 5 |

Table 1: Human genome, assembly GRCh38/hg38. Length of the autosomal chromosomes (nr of bases) and number of different bases in the two checks (see text).

| chr | #qualified | fraction qualified | #seg-ments | repeat masked | simple repeat | repeat | cds | introns | 3UTR | 5UTR | gene | all |
|---|---|---|---|---|---|---|---|---|---|---|---|---|
| hg38_chr1 | 229495936 | 0.922 | 248 | 0.37 | 0.005 | 0.518 | 0.014 | 0.602 | 0.023 | 0.009 | 0.635 | 1.0 |
| hg38_chr2 | 240353008 | 0.992 | 242 | 0.356 | 0.001 | 0.493 | 0.01 | 0.593 | 0.016 | 0.007 | 0.616 | 1.0 |
| hg38_chr3 | 197853613 | 0.998 | 198 | 0.371 | 0.033 | 0.514 | 0.009 | 0.623 | 0.016 | 0.006 | 0.645 | 1.0 |
| hg38_chr4 | 189591912 | 0.997 | 190 | 0.382 | 0.029 | 0.518 | 0.007 | 0.546 | 0.013 | 0.005 | 0.564 | 1.0 |
| hg38_chr5 | 180772919 | 0.996 | 181 | 0.377 | 0.002 | 0.515 | 0.008 | 0.56 | 0.015 | 0.006 | 0.581 | 1.0 |
| hg38_chr6 | 169326743 | 0.991 | 170 | 0.37 | 0.028 | 0.502 | 0.008 | 0.381 | 0.018 | 0.01 | 0.355 | 1.0 |
| hg38_chr7 | 158627095 | 0.995 | 159 | 0.387 | 0.051 | 0.514 | 0.008 | 0.404 | 0.018 | 0.01 | 0.373 | 1.0 |
| hg38_chr8 | 144685300 | 0.997 | 145 | 0.364 | 0.007 | 0.514 | 0.006 | 0.419 | 0.017 | 0.01 | 0.377 | 1.0 |
| hg38_chr9 | 121438540 | 0.877 | 138 | 0.372 | 0.044 | 0.518 | 0.009 | 0.378 | 0.019 | 0.011 | 0.349 | 1.0 |
| hg38_chr10 | 132497298 | 0.99 | 133 | 0.364 | 0.003 | 0.5 | 0.008 | 0.388 | 0.018 | 0.01 | 0.358 | 1.0 |
| hg38_chr11 | 134450920 | 0.995 | 135 | 0.37 | 0.052 | 0.527 | 0.014 | 0.583 | 0.021 | 0.01 | 0.614 | 1.0 |
| hg38_chr12 | 132861272 | 0.997 | 133 | 0.388 | 0.003 | 0.53 | 0.012 | 0.604 | 0.024 | 0.008 | 0.635 | 1.0 |
| hg38_chr13 | 97620716 | 0.854 | 98 | 0.368 | 0.04 | 0.493 | 0.005 | 0.312 | 0.015 | 0.008 | 0.277 | 1.0 |
| hg38_chr14 | 90558669 | 0.846 | 91 | 0.38 | 0.045 | 0.517 | 0.009 | 0.444 | 0.023 | 0.013 | 0.41 | 1.0 |
| hg38_chr15 | 83653532 | 0.82 | 84 | 0.373 | 0.053 | 0.515 | 0.011 | 0.464 | 0.028 | 0.016 | 0.433 | 1.0 |
| hg38_chr16 | 81569398 | 0.903 | 90 | 0.382 | 0.065 | 0.518 | 0.013 | 0.426 | 0.031 | 0.02 | 0.408 | 1.0 |
| hg38_chr17 | 82658588 | 0.993 | 83 | 0.403 | 0.008 | 0.521 | 0.022 | 0.625 | 0.031 | 0.013 | 0.67 | 1.0 |
| hg38_chr18 | 79812216 | 0.993 | 80 | 0.383 | 0.087 | 0.51 | 0.006 | 0.508 | 0.016 | 0.006 | 0.527 | 1.0 |
| hg38_chr19 | 57830142 | 0.987 | 58 | 0.467 | 0.109 | 0.599 | 0.034 | 0.607 | 0.043 | 0.017 | 0.669 | 1.0 |
| hg38_chr20 | 63576097 | 0.987 | 64 | 0.367 | 0.017 | 0.537 | 0.012 | 0.527 | 0.019 | 0.01 | 0.557 | 1.0 |
| hg38_chr21 | 39378764 | 0.843 | 41 | 0.391 | 0.022 | 0.518 | 0.007 | 0.515 | 0.016 | 0.01 | 0.537 | 1.0 |
| hg38_chr22 | 38832574 | 0.764 | 40 | 0.394 | 0.107 | 0.536 | 0.017 | 0.574 | 0.034 | 0.016 | 0.62 | 1.0 |

Table 2: Human, GRCh38/hg38. Statistics on input to the prediction. For explanation of the columns see the text.

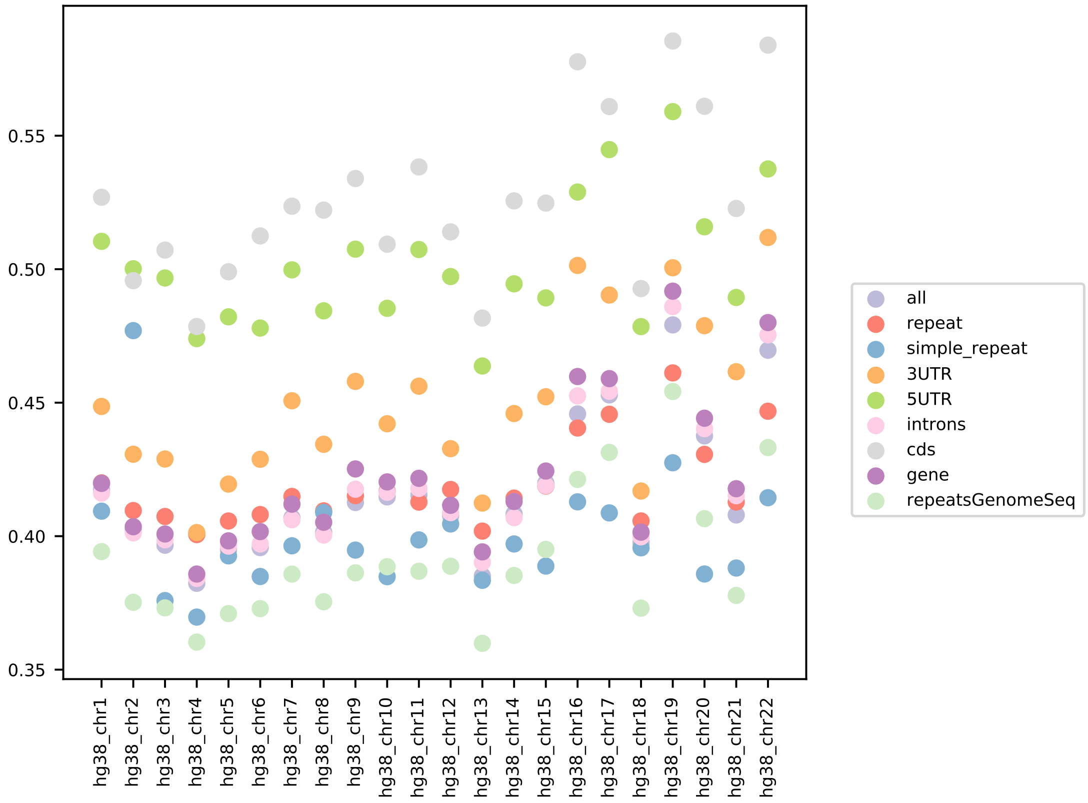


Figure 1: GC content per chromosome and annotation type for the human reference genome, assembly GRCh38/hg38.

| chr | all | repeat | simple repeat | repeat masked | cds | gene | introns | 3UTR | 5UTR |
|---|---|---|---|---|---|---|---|---|---|
| hg38_chr1 | 0.4173 | 0.42 | 0.4094 | 0.3942 | 0.5269 | 0.4198 | 0.4162 | 0.4486 | 0.5104 |
| hg38_chr2 | 0.4023 | 0.4096 | 0.477 | 0.3752 | 0.4957 | 0.4035 | 0.4012 | 0.4307 | 0.5002 |
| hg38_chr3 | 0.3966 | 0.4074 | 0.3759 | 0.3731 | 0.5072 | 0.4008 | 0.3986 | 0.4289 | 0.4967 |
| hg38_chr4 | 0.3823 | 0.4006 | 0.3697 | 0.3603 | 0.4785 | 0.3858 | 0.3842 | 0.4013 | 0.474 |
| hg38_chr5 | 0.3949 | 0.4057 | 0.3926 | 0.371 | 0.499 | 0.3982 | 0.3963 | 0.4195 | 0.4821 |
| hg38_chr6 | 0.3957 | 0.4081 | 0.3849 | 0.3729 | 0.5125 | 0.4017 | 0.3971 | 0.4288 | 0.4779 |
| hg38_chr7 | 0.4069 | 0.4149 | 0.3964 | 0.3858 | 0.5236 | 0.412 | 0.4061 | 0.4507 | 0.4998 |
| hg38_chr8 | 0.4015 | 0.4095 | 0.4089 | 0.3754 | 0.5221 | 0.4052 | 0.4004 | 0.4344 | 0.4844 |
| hg38_chr9 | 0.4126 | 0.4152 | 0.3948 | 0.3863 | 0.534 | 0.4252 | 0.4176 | 0.458 | 0.5075 |
| hg38_chr10 | 0.4147 | 0.4169 | 0.3848 | 0.3886 | 0.5094 | 0.4203 | 0.4164 | 0.4421 | 0.4853 |
| hg38_chr11 | 0.4154 | 0.4128 | 0.3986 | 0.3869 | 0.5383 | 0.4217 | 0.4179 | 0.4562 | 0.5074 |
| hg38_chr12 | 0.4076 | 0.4175 | 0.4046 | 0.3887 | 0.514 | 0.4116 | 0.4088 | 0.4328 | 0.4973 |
| hg38_chr13 | 0.3851 | 0.4019 | 0.3835 | 0.3599 | 0.4817 | 0.3941 | 0.3902 | 0.4124 | 0.4637 |
| hg38_chr14 | 0.4083 | 0.4142 | 0.3971 | 0.3853 | 0.5256 | 0.413 | 0.4069 | 0.4459 | 0.4945 |
| hg38_chr15 | 0.4199 | 0.4188 | 0.3888 | 0.3951 | 0.5248 | 0.4244 | 0.4189 | 0.4522 | 0.4893 |
| hg38_chr16 | 0.4458 | 0.4405 | 0.4129 | 0.4212 | 0.5777 | 0.4598 | 0.4525 | 0.5014 | 0.5289 |
| hg38_chr17 | 0.4529 | 0.4457 | 0.4087 | 0.4314 | 0.5609 | 0.459 | 0.4543 | 0.4903 | 0.5448 |
| hg38_chr18 | 0.3975 | 0.4057 | 0.3956 | 0.373 | 0.4927 | 0.4015 | 0.3998 | 0.4169 | 0.4785 |
| hg38_chr19 | 0.4791 | 0.4612 | 0.4275 | 0.4542 | 0.5855 | 0.4918 | 0.486 | 0.5005 | 0.559 |
| hg38_chr20 | 0.4376 | 0.4306 | 0.3858 | 0.4066 | 0.561 | 0.4442 | 0.4403 | 0.4788 | 0.5159 |
| hg38_chr21 | 0.4079 | 0.4128 | 0.3881 | 0.3778 | 0.5227 | 0.4178 | 0.4151 | 0.4616 | 0.4894 |
| hg38_chr22 | 0.4696 | 0.4468 | 0.4144 | 0.4332 | 0.5839 | 0.48 | 0.4754 | 0.5119 | 0.5375 |
| All | 0.4093 | 0.4152 | 0.3961 | 0.3854 | 0.527 | 0.4153 | 0.4113 | 0.448 | 0.5012 |

Table 3: Human, GRCh38/hg38. Fraction of GC content per annotation.

We carried out the two same tests for the mouse chromosomes that we did for the human and no differences were seen. The chromosome sequences used for the comparison were downloaded from [3] site http://hgdownload.soe.ucsc.edu/goldenPath/mm10/chromosomes/ (July, 2020).

Table 4 reports the ”statistics” for the input to the predictions and accuracy computations, just as Table 2 for the human case. As can be seen the fraction of qualified positions is high throughout.

### Zebrafish

The reference assembly GRCz11 (soft masked for showing repeats) was downloaded from [2], at site https://ftp.ncbi.nlm.nih.gov/genomes/refseq/vertebrate_other/Danio_rerio/all_assembly_versions/GCF_000002035.6_GRCz11/, the concrete file

| chr | length | #qualified | fraction qualified | #seg-ments | repeat masked | all |
|---|---|---|---|---|---|---|
| m38_chr1 | 195471971 | 191532614 | 0.98 | 195 | 0.457 | 1.0 |
| m38_chr2 | 182113224 | 178311077 | 0.979 | 182 | 0.426 | 1.0 |
| m38_chr3 | 160039680 | 156397328 | 0.977 | 160 | 0.465 | 1.0 |
| m38_chr4 | 156508116 | 151745996 | 0.97 | 156 | 0.463 | 1.0 |
| m38_chr5 | 151834684 | 147181940 | 0.969 | 151 | 0.442 | 1.0 |
| m38_chr6 | 149736546 | 145749271 | 0.973 | 149 | 0.447 | 1.0 |
| m38_chr7 | 145441459 | 141510898 | 0.973 | 145 | 0.476 | 1.0 |
| m38_chr8 | 129401213 | 125309169 | 0.968 | 129 | 0.427 | 1.0 |
| m38_chr9 | 124595110 | 120762514 | 0.969 | 124 | 0.435 | 1.0 |
| m38_chr10 | 130694993 | 126469919 | 0.968 | 130 | 0.445 | 1.0 |
| m38_chr11 | 122082543 | 118745645 | 0.973 | 122 | 0.413 | 1.0 |
| m38_chr12 | 120129022 | 116892948 | 0.973 | 120 | 0.44 | 1.0 |
| m38_chr13 | 120421639 | 116798804 | 0.97 | 120 | 0.444 | 1.0 |
| m38_chr14 | 124902244 | 120638761 | 0.966 | 124 | 0.441 | 1.0 |
| m38_chr15 | 104043685 | 100652515 | 0.967 | 104 | 0.433 | 1.0 |
| m38_chr16 | 98207768 | 94911106 | 0.966 | 98 | 0.436 | 1.0 |
| m38_chr17 | 94987271 | 90816441 | 0.956 | 94 | 0.448 | 1.0 |
| m38_chr18 | 90702639 | 86849245 | 0.958 | 90 | 0.435 | 1.0 |
| m38_chr19 | 61431566 | 57873940 | 0.942 | 61 | 0.419 | 1.0 |

Table 4: Mouse genome, assembly GRCm38/mm10. Statistics on input to the predictions.

being GCF_0000020- 35.6_GRCz11_genomic.fna.gz. As for the other genomes, the full (primary) sequence was subsequently split, resulting in 25 chromosomes used for predictions.

Table 5 reports the ”statistics” for the input to the predictions and accuracy computations, just as Table 2 for the human case. As can be seen the fraction of qualified positions is high throughout ($> 96$ %).

#### Fruit fly

The reference genome assembly was downloaded from [2], at https://ftp.ncbi.nlm-.nih.gov/genomes/genbank/invertebrate/Drosophila_melanogaster/all_assembly_versions, file: GCA_000001215.4_Release_6_plus_ISO1_MT and subsequently split so as to have the individual chromosome sequences.

Table 6 reports the ”statistics” for the input to the predictions and accuracy computations, just as Table 2 for the human case. As can be seen the fraction of qualified positions is high throughout ($> 97$ %) except for the shortest chromosome, chr4.

#### Yeast

The yeast genome assembly R64 was downloaded from [4] at site http://sgd-archive.yeastgenome.org/sequence/S288C_reference/genome_releases/. As with the human genome the file was split into files (sequences) for individual chromosomes.

We carried out the two same tests for the yeast chromosomes that we did for mouse and human; no differences were seen. The chromosome sequences used for the comparison were downloaded from [3] site https://hgdownload.soe.ucsc.edu/-goldenPath/sacCer3/chromosomes/ (July, 2020).

The simple repeats annotation sequence was downloaded from [3].

Table 7 reports the ”statistics” for the input to the predictions and accuracy computations, just as Table 2 for the human case. As can be seen, with one exception (chr6) the fraction of qualified positions is high throughout.

### Test of sampling in training procedure

This section is dedicated to a test of the uniformity of the sampling used in the training of the neural networks. Since the sampling was not used for the pre-

| chr | length | #qualified | fraction qualified | #seg-ments | repeat masked | all |
|---|---|---|---|---|---|---|
| GRCz11_chr1 | 59578282 | 58886847 | 0.988 | 59 | 0.473 | 1.0 |
| GRCz11_chr2 | 59640629 | 58842337 | 0.987 | 59 | 0.494 | 1.0 |
| GRCz11_chr3 | 62628489 | 61876776 | 0.988 | 62 | 0.479 | 1.0 |
| GRCz11_chr4 | 78093715 | 75126176 | 0.962 | 78 | 0.525 | 1.0 |
| GRCz11_chr5 | 72500376 | 71826576 | 0.991 | 72 | 0.493 | 1.0 |
| GRCz11_chr6 | 60270059 | 59851582 | 0.993 | 60 | 0.493 | 1.0 |
| GRCz11_chr7 | 74282399 | 73827529 | 0.994 | 74 | 0.477 | 1.0 |
| GRCz11_chr8 | 54304671 | 53908128 | 0.993 | 54 | 0.494 | 1.0 |
| GRCz11_chr9 | 56459846 | 55874391 | 0.99 | 56 | 0.483 | 1.0 |
| GRCz11_chr10 | 45420867 | 44904680 | 0.989 | 45 | 0.491 | 1.0 |
| GRCz11_chr11 | 45484837 | 44869208 | 0.986 | 45 | 0.488 | 1.0 |
| GRCz11_chr12 | 49182954 | 48818150 | 0.993 | 49 | 0.49 | 1.0 |
| GRCz11_chr13 | 52186027 | 51879672 | 0.994 | 52 | 0.488 | 1.0 |
| GRCz11_chr14 | 52660232 | 51895223 | 0.985 | 52 | 0.486 | 1.0 |
| GRCz11_chr15 | 48040578 | 47860510 | 0.996 | 48 | 0.492 | 1.0 |
| GRCz11_chr16 | 55266484 | 54883113 | 0.993 | 55 | 0.497 | 1.0 |
| GRCz11_chr17 | 53461100 | 52857727 | 0.989 | 53 | 0.491 | 1.0 |
| GRCz11_chr18 | 51023478 | 50812317 | 0.996 | 51 | 0.488 | 1.0 |
| GRCz11_chr19 | 48449771 | 47888635 | 0.988 | 48 | 0.485 | 1.0 |
| GRCz11_chr20 | 55201332 | 54860117 | 0.994 | 55 | 0.494 | 1.0 |
| GRCz11_chr21 | 45934066 | 44874023 | 0.977 | 45 | 0.494 | 1.0 |
| GRCz11_chr22 | 39133080 | 38867105 | 0.993 | 39 | 0.475 | 1.0 |
| GRCz11_chr23 | 46223584 | 45868239 | 0.992 | 46 | 0.488 | 1.0 |
| GRCz11_chr24 | 42172926 | 41871251 | 0.993 | 42 | 0.485 | 1.0 |
| GRCz11_chr25 | 37502051 | 36875129 | 0.983 | 37 | 0.494 | 1.0 |

Table 5: Zebrafish, GRCz11. Statistics on input to the predictions.

| **chr** | **#qualified** | **fraction qualified** | **#seg-ments** | **all** |
|---|---|---|---|---|
| r6.18_chrX | 22935030 | 0.974 | 23 | 1.0 |
| r6.18_chr2L | 32999550 | 0.99 | 33 | 1.0 |
| r6.18_chr2R | 24992450 | 0.988 | 25 | 1.0 |
| r6.18_chr3L | 27880890 | 0.992 | 28 | 1.0 |
| r6.18_chr3R | 31972678 | 0.997 | 32 | 1.0 |
| r6.18_chr4 | 999950 | 0.742 | 1 | 1.0 |

Table 6: Fruit fly, r6.18. Statistics on input to the predictions.

| **chr** | **length** | **#qualified** | **fraction qualified** | **#seg-ments** | **simple repeat** | **all** |
|---|---|---|---|---|---|---|
| R64_chr1 | 230218 | 199950 | 0.869 | 2 | 0.017 | 1.0 |
| R64_chr2 | 813184 | 799950 | 0.984 | 8 | 0.005 | 1.0 |
| R64_chr3 | 316620 | 299950 | 0.947 | 3 | 0.008 | 1.0 |
| R64_chr4 | 1531933 | 1499950 | 0.979 | 15 | na | 1.0 |
| R64_chr5 | 576874 | 499950 | 0.867 | 5 | na | 1.0 |
| R64_chr6 | 270161 | 199950 | 0.74 | 2 | na | 1.0 |
| R64_chr7 | 1090940 | 999950 | 0.917 | 10 | na | 1.0 |
| R64_chr8 | 562643 | 499950 | 0.889 | 5 | na | 1.0 |
| R64_chr9 | 439888 | 399950 | 0.909 | 4 | na | 1.0 |
| R64_chr10 | 745751 | 699950 | 0.939 | 7 | na | 1.0 |
| R64_chr11 | 666816 | 599950 | 0.9 | 6 | na | 1.0 |
| R64_chr12 | 1078177 | 999950 | 0.927 | 10 | na | 1.0 |
| R64_chr13 | 924431 | 899950 | 0.974 | 9 | na | 1.0 |
| R64_chr14 | 784333 | 699950 | 0.892 | 7 | na | 1.0 |
| R64_chr15 | 1091291 | 999950 | 0.916 | 10 | na | 1.0 |
| R64_chr16 | 948066 | 899950 | 0.949 | 9 | na | 1.0 |

Table 7: Yeast, R64. Statistics on input to the predictions.

dictions, this is to some extent only a ”nice-to-know”: were the sampling non-uniform over the genome only the training of the models could be harmed.

The test was done as follows. As explained in Supplementary Methods the training of the models were done in series of ”repeats” (”rounds” or ”big epochs”); each repeat consisted of 100 epochs, each in turn consisting in 100 steps of training batches of size 500. Thus each repeat uses 5 million samples. At the end of each repeat a validation was run based on 1 million samples. The training and validation samples were drawn from 4:1 division of the genomic sequence.

To test that this sampling was uniform, a full training sessions of 200 repeats was carried out, recording at every 10 repeats how many times each position was sampled (both for the training and for the validation). To reveal the uniformity the genome sequence was partitioned into adjacent windows of a set length. Two figures were then computed for every 10 repeats, based on all samples accumulated up to that repeat number:

1. Average occupancy: the average number of samples in the windows over the genome sequence

2. the standard deviation in the same set of occupancy numbers

This was carried out for two window sizes: 100000 and 1 million bases. Figure 2 below shows these averages as a function of the repeat number (one average every 10 repeats) with bars at each measurement indication 10 times the standard deviation (for the sake of visibility). Clearly, the sampling appears to be uniform.

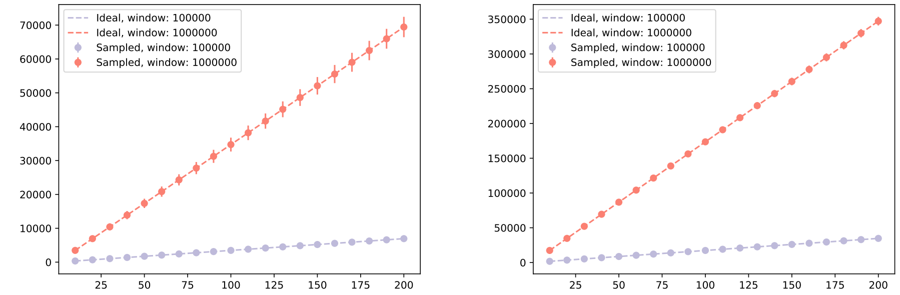


Figure 2: Results of sampling test. The plot to the left shows the results for the validation samples, the one to the right the training samples. Dots represent the average number of samples (given window size) and the bars show 10 standard deviations in the set of sampling occupancy numbers per window (on which the averages are had).
