## Supplementary I for "Prediction of DNA from context using neural networks"

### Supplementary I: Prediction of DNA from context using neural networks. Tables and additional plots.

May 13, 2021

### Human genome

| chr/model | LSTM1 | LSTM4S | LSTM4 | LSTM11 | mouseLSTM4 |
|---|---|---|---|---|---|
| hg38_chr1 | 0.5299 | 0.5204 | 0.519 | 0.5087 | 0.4347 |
| hg38_chr2 | 0.5163 | 0.5055 | 0.5056 | 0.4954 | 0.4278 |
| hg38_chr3 | 0.5224 | 0.5128 | 0.5114 | 0.4997 | 0.4258 |
| hg38_chr4 | 0.5218 | 0.5107 | 0.5106 | 0.4987 | 0.4245 |
| hg38_chr5 | 0.5267 | 0.5168 | 0.5155 | 0.5042 | 0.4251 |
| hg38_chr6 | 0.5193 | 0.5088 | 0.509 | 0.4978 | 0.4272 |
| hg38_chr7 | 0.5347 | 0.5257 | 0.5246 | 0.5145 | 0.4355 |
| hg38_chr8 | 0.5222 | 0.5097 | 0.5112 | 0.4998 | 0.4264 |
| hg38_chr9 | 0.5316 | 0.5218 | 0.5205 | 0.5094 | 0.4327 |
| hg38_chr10 | 0.5272 | 0.5161 | 0.5168 | 0.5066 | 0.4331 |
| hg38_chr11 | 0.5353 | 0.5252 | 0.5238 | 0.5127 | 0.4283 |
| hg38_chr12 | 0.5366 | 0.5248 | 0.5258 | 0.5146 | 0.4341 |
| hg38_chr13 | 0.5161 | 0.5064 | 0.5056 | 0.4945 | 0.4237 |
| hg38_chr14 | 0.533 | 0.5226 | 0.522 | 0.5107 | 0.4315 |
| hg38_chr15 | 0.5338 | 0.5262 | 0.5241 | 0.5137 | 0.4351 |
| hg38_chr16 | 0.5496 | 0.539 | 0.5397 | 0.5305 | 0.4484 |
| hg38_chr17 | 0.5641 | 0.5572 | 0.5556 | 0.5457 | 0.4557 |
| hg38_chr18 | 0.5356 | 0.519 | 0.5258 | 0.5148 | 0.4217 |
| hg38_chr19 | 0.6018 | 0.5962 | 0.594 | 0.5868 | 0.4811 |
| hg38_chr20 | 0.5414 | 0.5286 | 0.5313 | 0.5208 | 0.4357 |
| hg38_chr21 | 0.5399 | 0.5307 | 0.5297 | 0.5181 | 0.4276 |
| hg38_chr22 | 0.572 | 0.562 | 0.5619 | 0.5519 | 0.4553 |

Table 1: Accuracy of the predictions of the five models indicated per chromosome in human reference genome GRCh38 (hg38).

| anno/model | LSTM1 | LSTM4S | LSTM4 | LSTM11 | mouseLSTM4 |
|---|---|---|---|---|---|
| all | 0.5312 | 0.5209 | 0.5206 | 0.5098 | 0.4321 |
| repeat | 0.6542 | 0.6375 | 0.637 | 0.615 | 0.4692 |
| simple repeat | 0.8546 | 0.8389 | 0.8508 | 0.8272 | 0.4968 |
| 3UTR | 0.4461 | 0.4401 | 0.4396 | 0.4373 | 0.412 |
| 5UTR | 0.4383 | 0.4319 | 0.4316 | 0.4291 | 0.4066 |
| introns | 0.5195 | 0.51 | 0.5091 | 0.4998 | 0.4356 |
| cds | 0.3896 | 0.3856 | 0.3858 | 0.3873 | 0.3847 |
| gene | 0.5146 | 0.5052 | 0.5044 | 0.4956 | 0.434 |
| repeatsGenomeSeq | 0.7504 | 0.7414 | 0.7409 | 0.7204 | 0.5389 |

Table 2: Accuracy of the predictions of the five models indicated per annotation in human reference genome GRCh38 (hg38).

| chr | all | repeats | simple repeats | repeats masked | cds | gene | introns | 3UTR | 5UTR |
|---|---|---|---|---|---|---|---|---|---|
| hg38_chr1 | 0.5299 | 0.6501 | 0.795 | 0.7497 | 0.3897 | 0.5167 | 0.5218 | 0.4448 | 0.4424 |
| hg38_chr2 | 0.5163 | 0.6375 | 0.764 | 0.7335 | 0.3793 | 0.5069 | 0.5102 | 0.4431 | 0.4325 |
| hg38_chr3 | 0.5224 | 0.6412 | 0.8349 | 0.7379 | 0.3828 | 0.5094 | 0.5125 | 0.4387 | 0.438 |
| hg38_chr4 | 0.5218 | 0.6361 | 0.8132 | 0.7276 | 0.3727 | 0.513 | 0.5161 | 0.4422 | 0.4314 |
| hg38_chr5 | 0.5267 | 0.6484 | 0.9212 | 0.7426 | 0.3793 | 0.5115 | 0.5149 | 0.4328 | 0.4301 |
| hg38_chr6 | 0.5193 | 0.6407 | 0.7976 | 0.7334 | 0.3849 | 0.5042 | 0.5116 | 0.4426 | 0.4257 |
| hg38_chr7 | 0.5347 | 0.6607 | 0.8424 | 0.7522 | 0.3905 | 0.512 | 0.5184 | 0.438 | 0.4302 |
| hg38_chr8 | 0.5222 | 0.6396 | 0.9364 | 0.7397 | 0.3885 | 0.5056 | 0.5114 | 0.4401 | 0.4328 |
| hg38_chr9 | 0.5316 | 0.6536 | 0.8551 | 0.7517 | 0.3932 | 0.5123 | 0.5197 | 0.4385 | 0.4355 |
| hg38_chr10 | 0.5272 | 0.6541 | 0.9247 | 0.7497 | 0.3822 | 0.5101 | 0.5175 | 0.4401 | 0.4253 |
| hg38_chr11 | 0.5353 | 0.656 | 0.871 | 0.7598 | 0.3941 | 0.5135 | 0.5184 | 0.4428 | 0.4418 |
| hg38_chr12 | 0.5366 | 0.6587 | 0.8815 | 0.7547 | 0.3849 | 0.5185 | 0.5231 | 0.4526 | 0.4401 |
| hg38_chr13 | 0.5161 | 0.6364 | 0.8393 | 0.7257 | 0.378 | 0.4984 | 0.5041 | 0.4308 | 0.4308 |
| hg38_chr14 | 0.533 | 0.6572 | 0.8555 | 0.7521 | 0.3902 | 0.5157 | 0.5233 | 0.4438 | 0.4355 |
| hg38_chr15 | 0.5338 | 0.6608 | 0.8613 | 0.7606 | 0.3938 | 0.5174 | 0.5252 | 0.4483 | 0.4386 |
| hg38_chr16 | 0.5496 | 0.6802 | 0.8498 | 0.7795 | 0.4069 | 0.533 | 0.5413 | 0.4595 | 0.4537 |
| hg38_chr17 | 0.5641 | 0.7063 | 0.9502 | 0.7956 | 0.3997 | 0.5367 | 0.5443 | 0.4541 | 0.4516 |
| hg38_chr18 | 0.5356 | 0.6686 | 0.9122 | 0.7631 | 0.3746 | 0.4988 | 0.5016 | 0.4398 | 0.4421 |
| hg38_chr19 | 0.6018 | 0.7224 | 0.8437 | 0.8103 | 0.4092 | 0.5717 | 0.5844 | 0.4863 | 0.46 |
| hg38_chr20 | 0.5414 | 0.6607 | 0.9244 | 0.7752 | 0.398 | 0.5129 | 0.5175 | 0.4483 | 0.4511 |
| hg38_chr21 | 0.5399 | 0.6665 | 0.9667 | 0.7568 | 0.3851 | 0.503 | 0.5065 | 0.4416 | 0.437 |
| hg38_chr22 | 0.572 | 0.7035 | 0.8626 | 0.8036 | 0.4057 | 0.5367 | 0.5438 | 0.4726 | 0.4624 |
| All | 0.5312 | 0.6542 | 0.8546 | 0.7504 | 0.3896 | 0.5146 | 0.5195 | 0.4461 | 0.4383 |

Table 3: Accuracy of the predictions of LSTM1 per chromosome and annotation in human reference genome GRCh38 (hg38).

| chromo | all | repeats | simple repeats | repeats masked | cds | gene | introns | 3UTR | 5UTR |
|---|---|---|---|---|---|---|---|---|---|
| hg38_chr1 | 0.519 | 0.6325 | 0.7811 | 0.7399 | 0.3857 | 0.5063 | 0.5111 | 0.4385 | 0.4359 |
| hg38_chr2 | 0.5056 | 0.6193 | 0.7548 | 0.7231 | 0.3761 | 0.4967 | 0.4999 | 0.4368 | 0.4254 |
| hg38_chr3 | 0.5114 | 0.623 | 0.8337 | 0.7276 | 0.3788 | 0.4987 | 0.5016 | 0.4327 | 0.4306 |
| hg38_chr4 | 0.5106 | 0.6176 | 0.8116 | 0.7169 | 0.3694 | 0.5022 | 0.5051 | 0.4357 | 0.4248 |
| hg38_chr5 | 0.5155 | 0.6299 | 0.9179 | 0.7319 | 0.3763 | 0.5007 | 0.5039 | 0.4265 | 0.4222 |
| hg38_chr6 | 0.509 | 0.6234 | 0.8007 | 0.7236 | 0.382 | 0.4946 | 0.5016 | 0.4358 | 0.4192 |
| hg38_chr7 | 0.5246 | 0.6442 | 0.8353 | 0.7428 | 0.3859 | 0.5026 | 0.5086 | 0.4323 | 0.4246 |
| hg38_chr8 | 0.5112 | 0.6215 | 0.9432 | 0.7299 | 0.3858 | 0.4952 | 0.5007 | 0.4332 | 0.4256 |
| hg38_chr9 | 0.5205 | 0.6355 | 0.8474 | 0.7415 | 0.3896 | 0.5021 | 0.5091 | 0.4323 | 0.4286 |
| hg38_chr10 | 0.5168 | 0.6369 | 0.9299 | 0.7407 | 0.3789 | 0.5003 | 0.5074 | 0.4337 | 0.4194 |
| hg38_chr11 | 0.5238 | 0.6376 | 0.8646 | 0.7498 | 0.3893 | 0.5024 | 0.5071 | 0.4356 | 0.4342 |
| hg38_chr12 | 0.5258 | 0.6415 | 0.8727 | 0.7452 | 0.3817 | 0.5083 | 0.5126 | 0.4454 | 0.4326 |
| hg38_chr13 | 0.5056 | 0.6185 | 0.8353 | 0.7154 | 0.3742 | 0.4885 | 0.4941 | 0.4243 | 0.4235 |
| hg38_chr14 | 0.522 | 0.6394 | 0.8455 | 0.7418 | 0.3864 | 0.5054 | 0.5126 | 0.4371 | 0.4294 |
| hg38_chr15 | 0.5241 | 0.6452 | 0.8676 | 0.753 | 0.3903 | 0.5075 | 0.5148 | 0.4418 | 0.4326 |
| hg38_chr16 | 0.5397 | 0.6649 | 0.8441 | 0.7723 | 0.403 | 0.524 | 0.5318 | 0.4532 | 0.4475 |
| hg38_chr17 | 0.5556 | 0.6938 | 0.9525 | 0.7899 | 0.3951 | 0.5281 | 0.5354 | 0.4479 | 0.445 |
| hg38_chr18 | 0.5258 | 0.6527 | 0.9112 | 0.7547 | 0.3718 | 0.4888 | 0.4914 | 0.4334 | 0.434 |
| hg38_chr19 | 0.594 | 0.7125 | 0.8387 | 0.805 | 0.4041 | 0.5637 | 0.5762 | 0.4789 | 0.4541 |
| hg38_chr20 | 0.5313 | 0.6452 | 0.9294 | 0.7678 | 0.3932 | 0.5028 | 0.5072 | 0.4415 | 0.4438 |
| hg38_chr21 | 0.5297 | 0.6502 | 0.9698 | 0.7474 | 0.3814 | 0.4929 | 0.4962 | 0.4357 | 0.4294 |
| hg38_chr22 | 0.5619 | 0.6882 | 0.8503 | 0.7956 | 0.402 | 0.5273 | 0.5341 | 0.4664 | 0.4552 |
| All | 0.5206 | 0.637 | 0.8508 | 0.7409 | 0.3858 | 0.5044 | 0.5091 | 0.4396 | 0.4316 |

Table 4: Accuracy of the predictions of LSTM4 per chromosome and annotation in human reference genome GRCh38.

| chromo | all | repeats | simple repeats | repeats masked | cds | gene | introns | 3UTR | 5UTR |
|---|---|---|---|---|---|---|---|---|---|
| hg38_chr1 | 0.5087 | 0.6114 | 0.7752 | 0.7207 | 0.3877 | 0.4973 | 0.5016 | 0.4365 | 0.433 |
| hg38_chr2 | 0.4954 | 0.5974 | 0.7464 | 0.7027 | 0.3771 | 0.4878 | 0.4906 | 0.4343 | 0.4227 |
| hg38_chr3 | 0.4997 | 0.5993 | 0.8041 | 0.7049 | 0.3806 | 0.4889 | 0.4915 | 0.4301 | 0.4271 |
| hg38_chr4 | 0.4987 | 0.5937 | 0.7879 | 0.6944 | 0.3701 | 0.4917 | 0.4944 | 0.4319 | 0.4206 |
| hg38_chr5 | 0.5042 | 0.607 | 0.902 | 0.7107 | 0.3771 | 0.4908 | 0.4937 | 0.4249 | 0.4184 |
| hg38_chr6 | 0.4978 | 0.6 | 0.7692 | 0.7016 | 0.3837 | 0.4859 | 0.4924 | 0.4328 | 0.4167 |
| hg38_chr7 | 0.5145 | 0.6236 | 0.8196 | 0.7239 | 0.3887 | 0.4946 | 0.4998 | 0.4311 | 0.4227 |
| hg38_chr8 | 0.4998 | 0.5986 | 0.8736 | 0.7076 | 0.3865 | 0.4862 | 0.4912 | 0.4303 | 0.4233 |
| hg38_chr9 | 0.5094 | 0.6129 | 0.8271 | 0.7205 | 0.3906 | 0.4937 | 0.4999 | 0.4298 | 0.4271 |
| hg38_chr10 | 0.5066 | 0.6154 | 0.8885 | 0.7208 | 0.3802 | 0.4923 | 0.4987 | 0.4312 | 0.4178 |
| hg38_chr11 | 0.5127 | 0.6152 | 0.8493 | 0.7289 | 0.3911 | 0.4929 | 0.4971 | 0.4334 | 0.4314 |
| hg38_chr12 | 0.5146 | 0.6194 | 0.8436 | 0.7247 | 0.3835 | 0.4991 | 0.503 | 0.4421 | 0.4305 |
| hg38_chr13 | 0.4945 | 0.5951 | 0.8085 | 0.6938 | 0.376 | 0.4797 | 0.4848 | 0.4211 | 0.4195 |
| hg38_chr14 | 0.5107 | 0.6163 | 0.8173 | 0.7198 | 0.388 | 0.4962 | 0.5027 | 0.4347 | 0.4273 |
| hg38_chr15 | 0.5137 | 0.6239 | 0.8352 | 0.7326 | 0.3907 | 0.4991 | 0.5058 | 0.4383 | 0.429 |
| hg38_chr16 | 0.5305 | 0.6462 | 0.828 | 0.7563 | 0.4044 | 0.517 | 0.524 | 0.4518 | 0.4458 |
| hg38_chr17 | 0.5457 | 0.673 | 0.8985 | 0.7705 | 0.397 | 0.5217 | 0.5286 | 0.4472 | 0.4437 |
| hg38_chr18 | 0.5148 | 0.6302 | 0.8845 | 0.7331 | 0.3724 | 0.4803 | 0.4827 | 0.4312 | 0.4304 |
| hg38_chr19 | 0.5868 | 0.6987 | 0.8281 | 0.7927 | 0.4061 | 0.5583 | 0.5702 | 0.4766 | 0.4522 |
| hg38_chr20 | 0.5208 | 0.6246 | 0.8945 | 0.7477 | 0.395 | 0.4951 | 0.499 | 0.4402 | 0.441 |
| hg38_chr21 | 0.5181 | 0.6266 | 0.9228 | 0.7245 | 0.382 | 0.4845 | 0.4875 | 0.4333 | 0.4264 |
| hg38_chr22 | 0.5519 | 0.6681 | 0.823 | 0.7765 | 0.4026 | 0.5204 | 0.5268 | 0.4641 | 0.453 |
| All | 0.5098 | 0.615 | 0.8272 | 0.7204 | 0.3873 | 0.4956 | 0.4998 | 0.4373 | 0.4291 |

Table 5: Accuracy of the predictions of LSTM11 per chromosome and annotation in human reference genome GRCh38.

| chromo | all | repeats | simple repeats | repeats masked | cds | gene | introns | 3UTR | 5UTR |
|---|---|---|---|---|---|---|---|---|---|
| hg38_chr1 | 0.4347 | 0.4742 | 0.5314 | 0.5474 | 0.3849 | 0.4348 | 0.4366 | 0.4122 | 0.4098 |
| hg38_chr2 | 0.4278 | 0.4653 | 0.4544 | 0.5388 | 0.3749 | 0.4286 | 0.4298 | 0.4088 | 0.402 |
| hg38_chr3 | 0.4258 | 0.4598 | 0.5285 | 0.531 | 0.3785 | 0.4262 | 0.4271 | 0.4073 | 0.4035 |
| hg38_chr4 | 0.4245 | 0.454 | 0.5513 | 0.5241 | 0.3692 | 0.4257 | 0.4267 | 0.4065 | 0.3971 |
| hg38_chr5 | 0.4251 | 0.4575 | 0.3977 | 0.5259 | 0.375 | 0.4263 | 0.4274 | 0.4052 | 0.3971 |
| hg38_chr6 | 0.4272 | 0.4634 | 0.5607 | 0.5345 | 0.3794 | 0.4274 | 0.43 | 0.407 | 0.3974 |
| hg38_chr7 | 0.4355 | 0.4755 | 0.5101 | 0.5429 | 0.3847 | 0.4352 | 0.4368 | 0.4098 | 0.404 |
| hg38_chr8 | 0.4264 | 0.4597 | 0.3849 | 0.5322 | 0.3835 | 0.4274 | 0.4289 | 0.406 | 0.4019 |
| hg38_chr9 | 0.4327 | 0.4696 | 0.5226 | 0.5405 | 0.3885 | 0.4348 | 0.4367 | 0.4085 | 0.4066 |
| hg38_chr10 | 0.4331 | 0.4743 | 0.3858 | 0.5453 | 0.3776 | 0.4341 | 0.4369 | 0.4081 | 0.3997 |
| hg38_chr11 | 0.4283 | 0.4602 | 0.4809 | 0.5292 | 0.3883 | 0.4299 | 0.4315 | 0.4093 | 0.4082 |
| hg38_chr12 | 0.4341 | 0.472 | 0.4058 | 0.5422 | 0.3802 | 0.4359 | 0.4376 | 0.4113 | 0.4056 |
| hg38_chr13 | 0.4237 | 0.4555 | 0.4924 | 0.5253 | 0.3728 | 0.425 | 0.4271 | 0.4006 | 0.3979 |
| hg38_chr14 | 0.4315 | 0.468 | 0.4977 | 0.5361 | 0.3844 | 0.4328 | 0.4355 | 0.4085 | 0.4058 |
| hg38_chr15 | 0.4351 | 0.4764 | 0.4908 | 0.5467 | 0.3888 | 0.4378 | 0.4403 | 0.4118 | 0.4055 |
| hg38_chr16 | 0.4484 | 0.4977 | 0.5183 | 0.5683 | 0.3997 | 0.4543 | 0.457 | 0.4247 | 0.4203 |
| hg38_chr17 | 0.4557 | 0.5075 | 0.3953 | 0.5702 | 0.3942 | 0.4583 | 0.4617 | 0.4206 | 0.4203 |
| hg38_chr18 | 0.4217 | 0.4514 | 0.4263 | 0.5117 | 0.372 | 0.4255 | 0.4265 | 0.4072 | 0.4037 |
| hg38_chr19 | 0.4811 | 0.5286 | 0.5106 | 0.5875 | 0.4053 | 0.4839 | 0.4903 | 0.439 | 0.4259 |
| hg38_chr20 | 0.4357 | 0.4717 | 0.3914 | 0.5437 | 0.3918 | 0.439 | 0.4407 | 0.4163 | 0.4125 |
| hg38_chr21 | 0.4276 | 0.4564 | 0.3638 | 0.5163 | 0.38 | 0.4306 | 0.432 | 0.4101 | 0.4043 |
| hg38_chr22 | 0.4553 | 0.4968 | 0.469 | 0.5587 | 0.3998 | 0.4591 | 0.4622 | 0.4327 | 0.4237 |
| All | 0.4321 | 0.4692 | 0.4968 | 0.5389 | 0.3847 | 0.434 | 0.4356 | 0.412 | 0.4066 |

Table 6: Accuracy of the predictions of mouseLSTM4 per chromosome and annotation in human reference genome GRCh38.

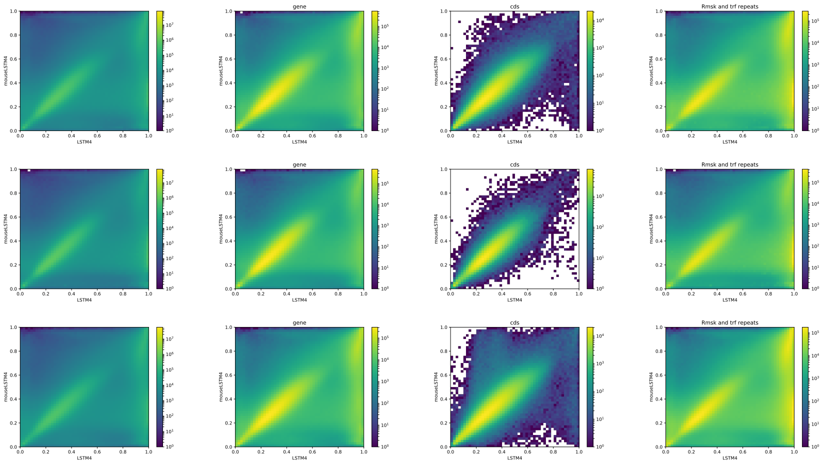


Figure 1: LSTM4 vs mouseLSTM4human. Scatter-plot of probabilities of reference bases in annotated parts of chromosomes of the human reference genome GRCh38 according to LSTM4 (x-axis) and mouseLSTM4 (y-axis). From the top: chromosome 17, 18, 19. Annotations from left to right: all positions, gene, cds, repeats.

| **Model1** | **Model2** | **Test value** | **p-value** |
|---|---|---|---|
| LSTM1 | k=3 central | 1877.6 | $< 10^{-20}$ |
| LSTM1 | k=4 central | 1646.4 | $< 10^{-20}$ |
| LSTM1 | k=5 central | 1339.9 | $< 10^{-20}$ |
| LSTM1 | Markov k=14 | 768.6 | $< 10^{-20}$ |

Table 7: Results of likelihood ratio tests. Model2 was used as base (”denominator”) in likelihood ratio test.

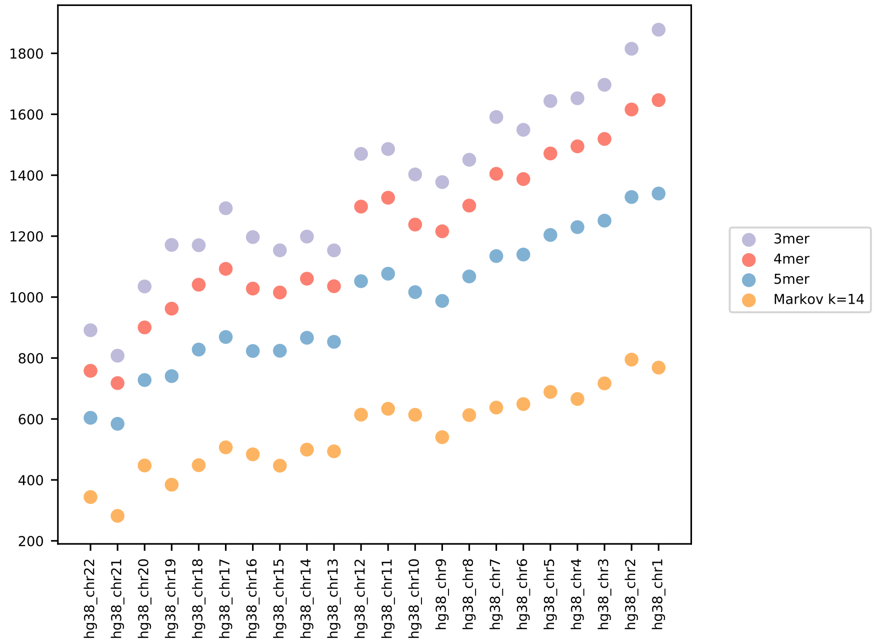


Figure 2: Likelihood ratio test figures per chromosome for LSTM1 vs simpler models as indicated. Test values increase with the size of the sample size (equivalently the chromosome size) supporting the rejection of the null hypotheses of equally performing models [1].

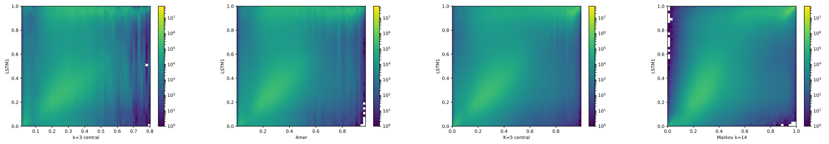

Figure 3: LSTM1 (x-axis) vs simpler models (y-axis) from left to right: central model for k=3,4,5 and Markov model, k=14. Each plot shows the density of the reference-base probabilities according to the named models. Please note that colors are not shared between the plots.

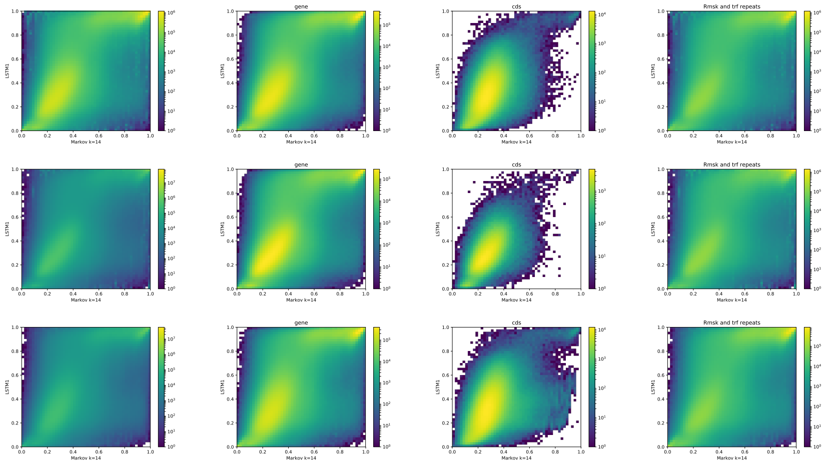


Figure 4: Markov k=14 vs. LSTM1. Scatter-plot of probabilities of reference bases in annotated parts of chromosomes of the human reference genome GRCh38 according to the Markov k=14 model(x-axis) and LSTM1 (y-axis). From the top: chromosome 17, 18, 19. Annotations from left to right: all positions, gene, cds, repeats.

### Other genomes

#### Yeast, S.cerevisae

| chr/annotation | all | simple repeats |
|---|---|---|
| R64_chr1 | 0.369 | 0.4239 |
| R64_chr2 | 0.372 | 0.4599 |
| R64_chr3 | 0.3719 | 0.4591 |
| R64_chr4 | 0.3744 | na |
| R64_chr5 | 0.3681 | na |
| R64_chr6 | 0.3705 | na |
| R64_chr7 | 0.3728 | na |
| R64_chr8 | 0.3704 | na |
| R64_chr9 | 0.3707 | na |
| R64_chr10 | 0.3705 | na |
| R64_chr11 | 0.3731 | na |
| R64_chr12 | 0.3704 | na |
| R64_chr13 | 0.3733 | na |
| R64_chr14 | 0.3693 | na |
| R64_chr15 | 0.372 | na |
| R64_chr16 | 0.3721 | na |
| All | 0.3718 | 0.4476 |

| chr/annotation | all | simple repeats |
|---|---|---|
| R64_chr1 | 0.4221 | 0.6612 |
| R64_chr2 | 0.416 | 0.6167 |
| R64_chr3 | 0.4152 | 0.6394 |
| R64_chr4 | 0.4209 | na |
| R64_chr5 | 0.4169 | na |
| R64_chr6 | 0.4194 | na |
| R64_chr7 | 0.4191 | na |
| R64_chr8 | 0.4077 | na |
| R64_chr9 | 0.4101 | na |
| R64_chr10 | 0.4152 | na |
| R64_chr11 | 0.4081 | na |
| R64_chr12 | 0.4201 | na |
| R64_chr13 | 0.4192 | na |
| R64_chr14 | 0.4133 | na |
| R64_chr15 | 0.4135 | na |
| R64_chr16 | 0.4192 | na |
| All | 0.4166 | 0.6373 |

Table 8: Accuracy of the predictions of LSTM41 (left) and LSTM4 (right) on yeast genome R64. LSTM41 is identical to LSTM4, but less trained (see Suppl. methods).

#### Fruit fly, D.melanogaster

| chr/annotation | all |
|---|---|
| r6.18_chrX | 0.4359 |
| r6.18_chr2L | 0.4149 |
| r6.18_chr2R | 0.4105 |
| r6.18_chr3L | 0.4165 |
| r6.18_chr3R | 0.4183 |
| r6.18_chr4 | 0.4157 |
| All | 0.4186 |

| chr/annotation | all |
|---|---|
| r6.18_chrX | 0.466 |
| r6.18_chr2L | 0.4468 |
| r6.18_chr2R | 0.4602 |
| r6.18_chr3L | 0.4604 |
| r6.18_chr3R | 0.4552 |
| r6.18_chr4 | 0.4443 |
| All | 0.4568 |

Table 9: Accuracy of the predictions of LSTM1 (left) and LSTM4 (right) on fruit fly genome dm6 (r6.18). LSTM41 is identical to LSTM4, but less trained (see Suppl. methods)

#### Zebrafish, D.rerio

| chr/annotation | all | repeats masked |
|---|---|---|
| GRCz11_chr1 | 0.5377 | 0.7019 |
| GRCz11_chr2 | 0.5463 | 0.7067 |
| GRCz11_chr3 | 0.5387 | 0.7004 |
| GRCz11_chr4 | 0.5212 | 0.6504 |
| GRCz11_chr5 | 0.5448 | 0.7043 |
| GRCz11_chr6 | 0.5448 | 0.704 |
| GRCz11_chr7 | 0.54 | 0.7066 |
| GRCz11_chr8 | 0.5454 | 0.7048 |
| GRCz11_chr9 | 0.5419 | 0.7053 |
| GRCz11_chr10 | 0.5471 | 0.7088 |
| GRCz11_chr11 | 0.5443 | 0.7054 |
| GRCz11_chr12 | 0.5454 | 0.7066 |
| GRCz11_chr13 | 0.5464 | 0.7098 |
| GRCz11_chr14 | 0.5427 | 0.7062 |
| GRCz11_chr15 | 0.5452 | 0.7052 |
| GRCz11_chr16 | 0.5483 | 0.7081 |
| GRCz11_chr17 | 0.5471 | 0.7098 |
| GRCz11_chr18 | 0.5448 | 0.7075 |
| GRCz11_chr19 | 0.5426 | 0.704 |
| GRCz11_chr20 | 0.5458 | 0.7052 |
| GRCz11_chr21 | 0.5459 | 0.7051 |
| GRCz11_chr22 | 0.5366 | 0.6993 |
| GRCz11_chr23 | 0.5455 | 0.7079 |
| GRCz11_chr24 | 0.545 | 0.7085 |
| GRCz11_chr25 | 0.5478 | 0.7089 |
| All | 0.5428 | 0.7024 |

Table 10: Accuracy of the predictions of LSTM4 on zebrafish genome GRCz11.

#### Mouse

| chr/annotation | all | repeats masked |
|---|---|---|
| m38_chr1 | 0.5136 | 0.6684 |
| m38_chr2 | 0.5034 | 0.6643 |
| m38_chr3 | 0.5188 | 0.6751 |
| m38_chr4 | 0.5157 | 0.6664 |
| m38_chr5 | 0.5105 | 0.6688 |
| m38_chr6 | 0.51 | 0.6662 |
| m38_chr7 | 0.5179 | 0.6653 |
| m38_chr8 | 0.5036 | 0.6653 |
| m38_chr9 | 0.5024 | 0.6573 |
| m38_chr10 | 0.5085 | 0.6661 |
| m38_chr11 | 0.4984 | 0.6588 |
| m38_chr12 | 0.5059 | 0.6618 |
| m38_chr13 | 0.5034 | 0.6573 |
| m38_chr14 | 0.5077 | 0.6664 |
| m38_chr15 | 0.5042 | 0.6618 |
| m38_chr16 | 0.5043 | 0.6607 |
| m38_chr17 | 0.5089 | 0.6617 |
| m38_chr18 | 0.5026 | 0.6603 |
| m38_chr19 | 0.4951 | 0.6526 |
| All | 0.5081 | 0.6644 |

Table 11: Accuracy of the predictions of mouseLSTM4 on mouse reference genome GRCm38 (mm10).
