## Supplementary IIc for "Prediction of DNA from context using neural networks"

### Supplementary IIc: Prediction of DNA from context using neural networks. Fourier, human genome, GC/AT content, low frequencies.

Christian Grønbæk, Yuhu Liang, Desmond Elliott, and Anders Krogh

February 6, 2021

This file contains the first of two sets of plots based on the Fourier analysis showing the L2-norm of the Fourier coefficients in a running window. This first set covers the frequency range from 200 to 45000 using a window length of 1000 (and a step size of 100); the second set covers the frequencies from 40000 to 140000 and used a window length of 5000 (and step size of 100) and is placed in a separate file.

### Fourier plots, GC/AT content, frequency range 200-45000

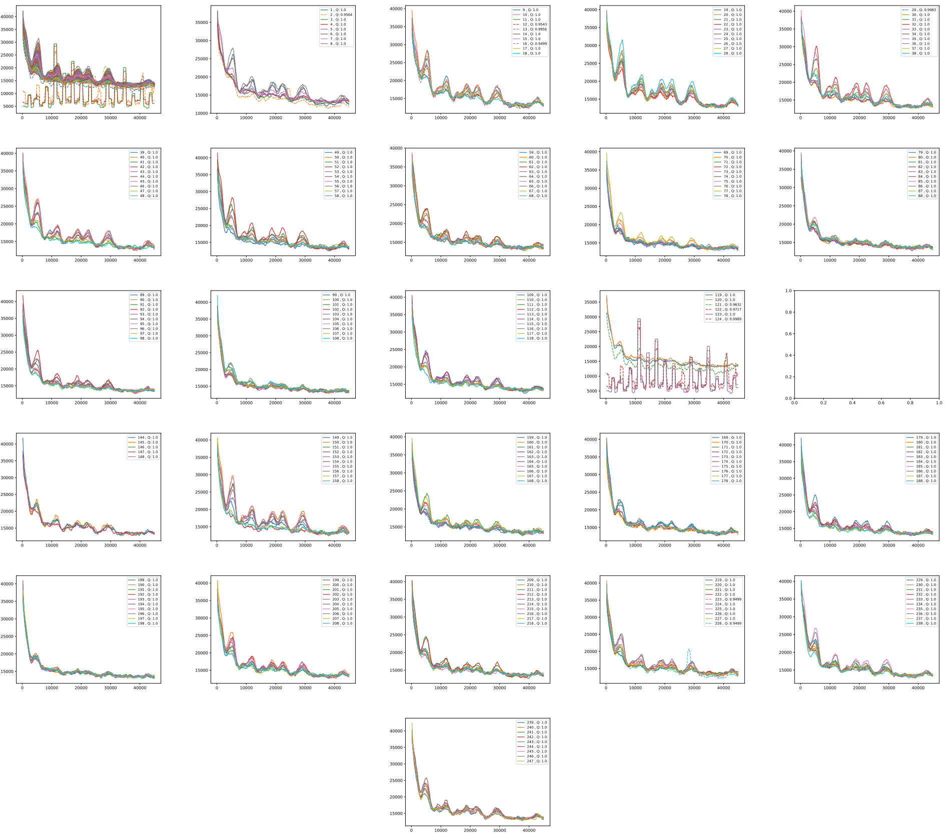

Figure 1: GC/AT content. Fouriers on GC/AT content arrays for chromosome: hg38_chr1. The genome string is divided in adjacent segments of 1Mb; several adjacent segments are covered in each plot, segment numbers are indicated in the legend. First plot (upper left) shows the results for all segments in this chromosome.

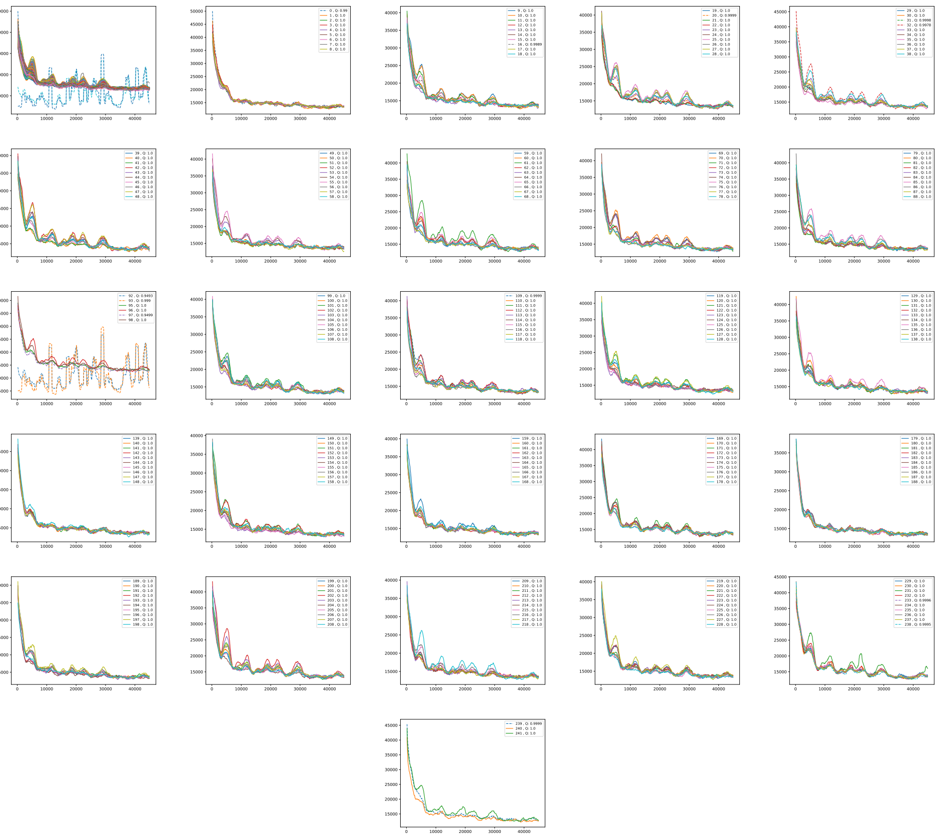

Figure 2: GC/AT content. Fouriers on GC/AT content arrays for chromosome: hg38_chr2. The genome string is divided in adjacent segments of 1Mb; several adjacent segments are covered in each plot, segment numbers are indicated in the legend. First plot (upper left) shows the results for all segments in this chromosome.

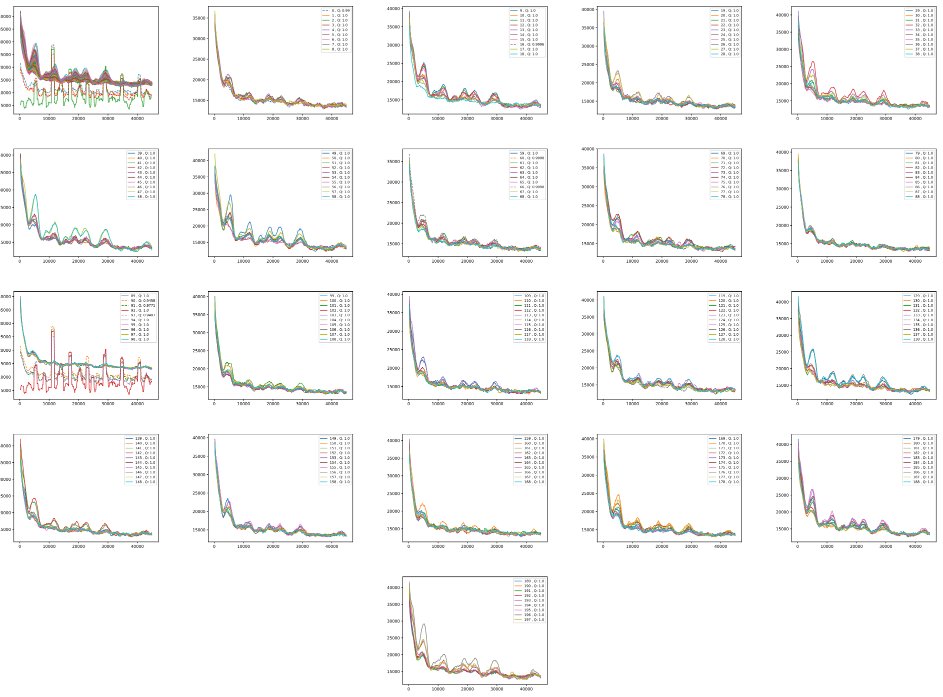

Figure 3: GC/AT content. Fouriers on GC/AT content arrays for chromosome: hg38_chr3. The genome string is divided in adjacent segments of 1Mb; several adjacent segments are covered in each plot, segment numbers are indicated in the legend. First plot (upper left) shows the results for all segments in this chromosome.

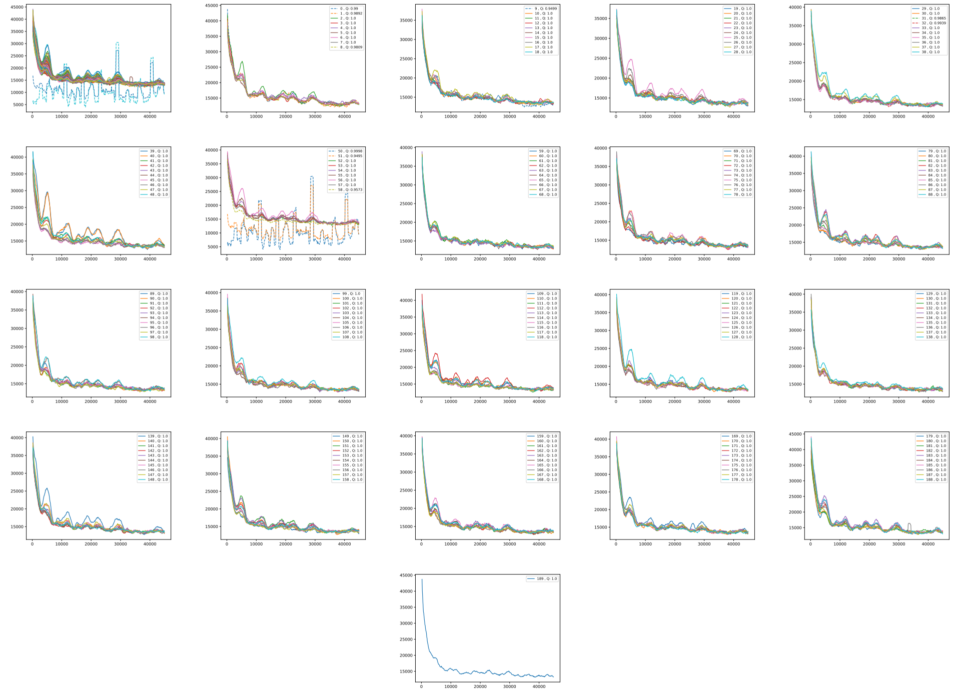

Figure 4: GC/AT content. Fouriers on GC/AT content arrays for chromosome: hg38_chr4. The genome string is divided in adjacent segments of 1Mb; several adjacent segments are covered in each plot, segment numbers are indicated in the legend. First plot (upper left) shows the results for all segments in this chromosome.

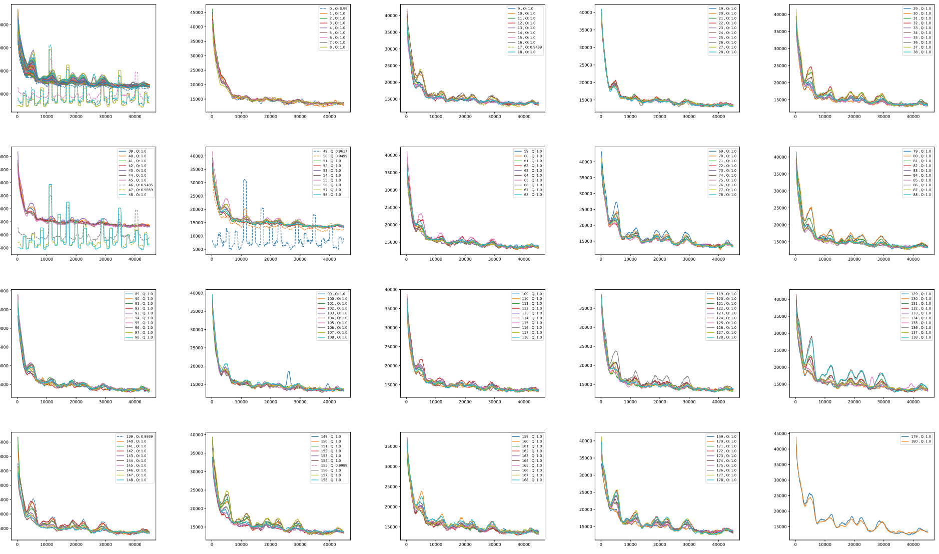

Figure 5: GC/AT content. Fouriers on GC/AT content arrays for chromosome: hg38_chr5. The genome string is divided in adjacent segments of 1Mb; several adjacent segments are covered in each plot, segment numbers are indicated in the legend. First plot (upper left) shows the results for all segments in this chromosome.

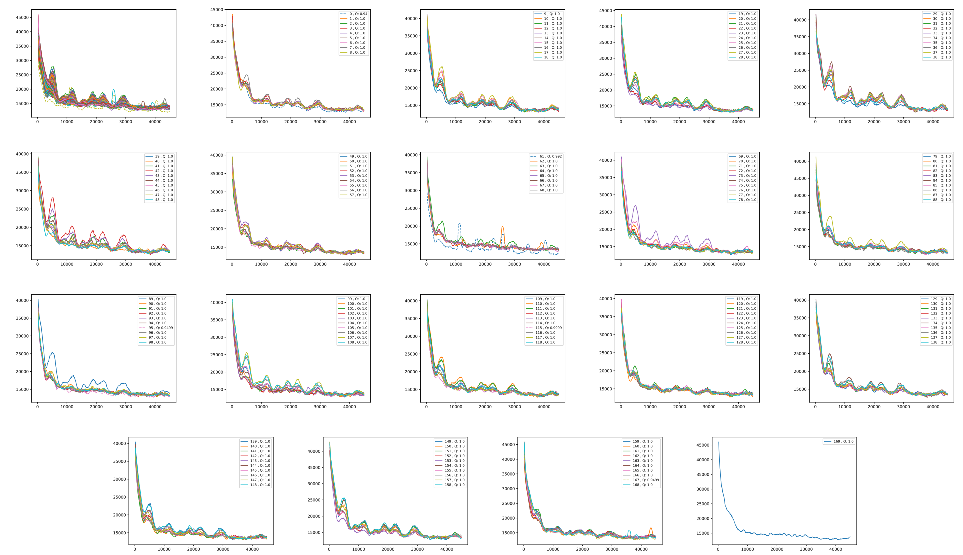

Figure 6: GC/AT content. Fouriers on GC/AT content arrays for chromosome: hg38_chr6. The genome string is divided in adjacent segments of 1Mb; several adjacent segments are covered in each plot, segment numbers are indicated in the legend. First plot (upper left) shows the results for all segments in this chromosome.

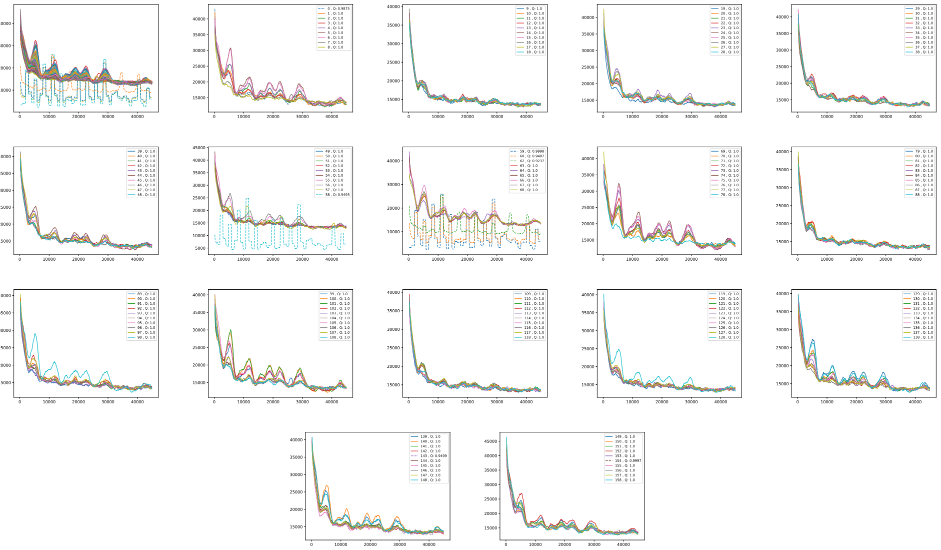

Figure 7: GC/AT content. Fouriers on GC/AT content arrays for chromosome: hg38_chr7. The genome string is divided in adjacent segments of 1Mb; several adjacent segments are covered in each plot, segment numbers are indicated in the legend. First plot (upper left) shows the results for all segments in this chromosome.

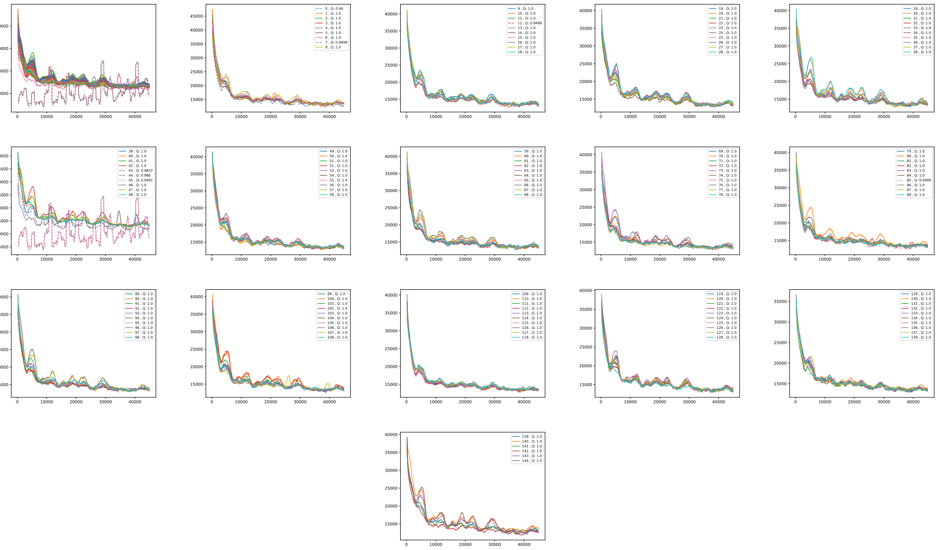

Figure 8: GC/AT content. Fouriers on GC/AT content arrays for chromosome: hg38_chr8. The genome string is divided in adjacent segments of 1Mb; several adjacent segments are covered in each plot, segment numbers are indicated in the legend. First plot (upper left) shows the results for all segments in this chromosome.

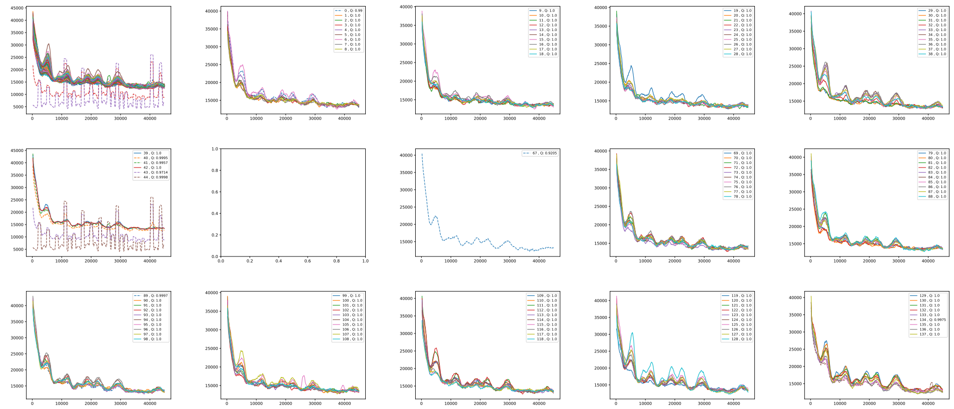

Figure 9: GC/AT content. Fouriers on GC/AT content arrays for chromosome: hg38_chr9. The genome string is divided in adjacent segments of 1Mb; several adjacent segments are covered in each plot, segment numbers are indicated in the legend. First plot (upper left) shows the results for all segments in this chromosome.

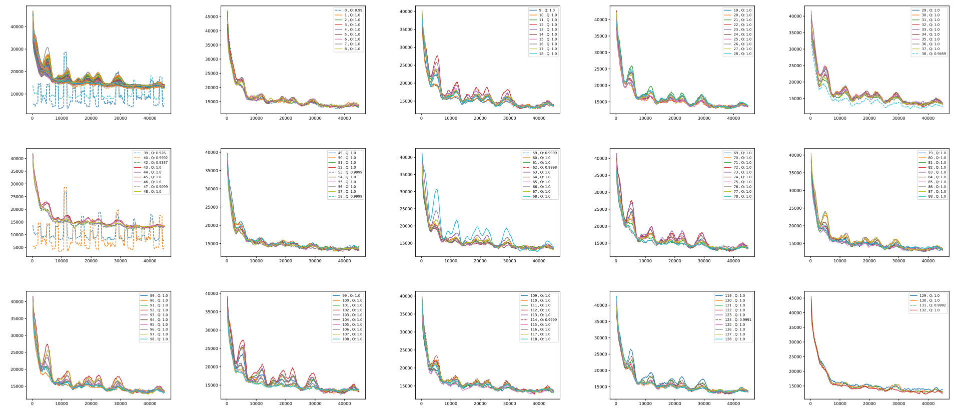

Figure 10: GC/AT content. Fouriers on GC/AT content arrays for chromosome: hg38_chr10. The genome string is divided in adjacent segments of 1Mb; several adjacent segments are covered in each plot, segment numbers are indicated in the legend. First plot (upper left) shows the results for all segments in this chromosome.

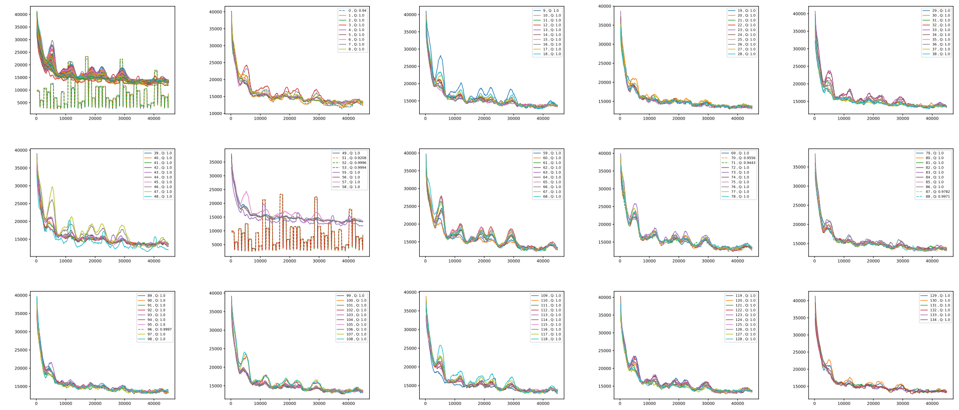

Figure 11: GC/AT content. Fouriers on GC/AT content arrays for chromosome: hg38_chr11. The genome string is divided in adjacent segments of 1Mb; several adjacent segments are covered in each plot, segment numbers are indicated in the legend. First plot (upper left) shows the results for all segments in this chromosome.

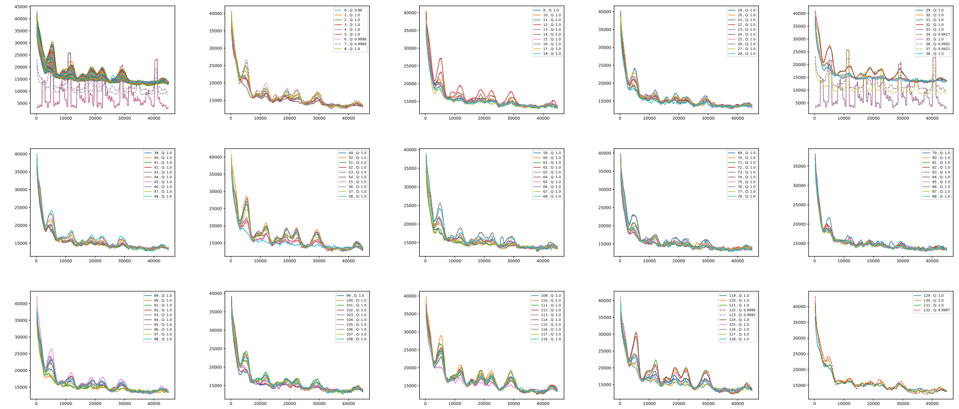

Figure 12: GC/AT content. Fouriers on GC/AT content arrays for chromosome: hg38_chr12. The genome string is divided in adjacent segments of 1Mb; several adjacent segments are covered in each plot, segment numbers are indicated in the legend. First plot (upper left) shows the results for all segments in this chromosome.

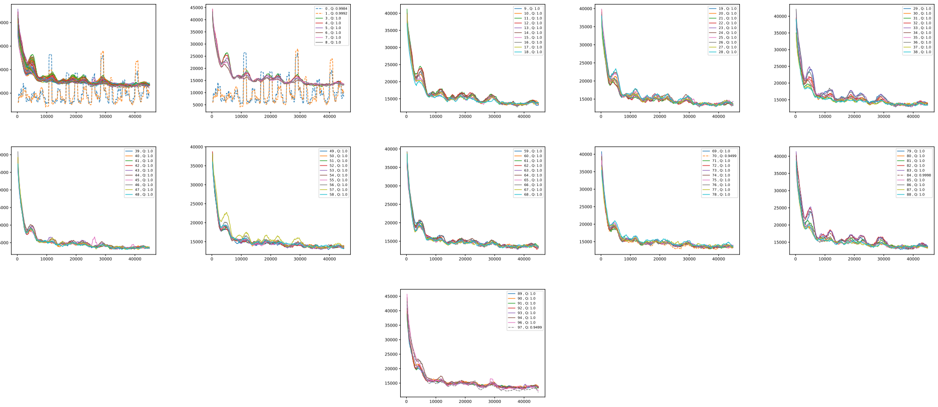

Figure 13: GC/AT content. Fouriers on GC/AT content arrays for chromosome: hg38_chr13. The genome string is divided in adjacent segments of 1Mb; several adjacent segments are covered in each plot, segment numbers are indicated in the legend. First plot (upper left) shows the results for all segments in this chromosome.

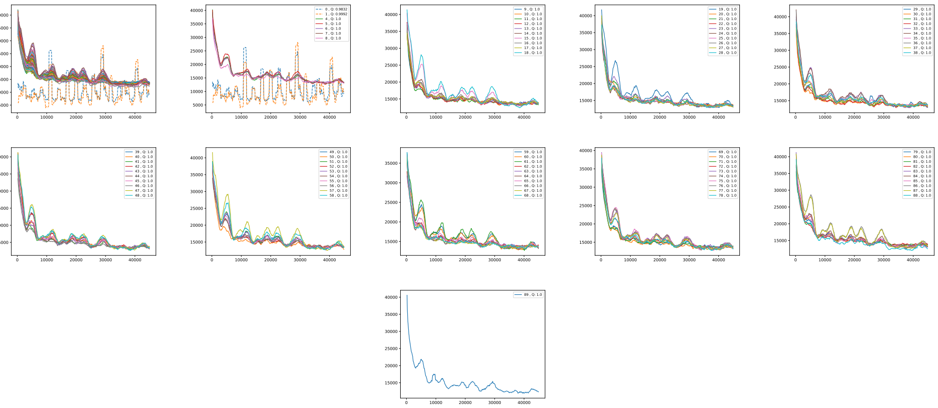

Figure 14: GC/AT content. Fouriers on GC/AT content arrays for chromosome: hg38_chr14. The genome string is divided in adjacent segments of 1Mb; several adjacent segments are covered in each plot, segment numbers are indicated in the legend. First plot (upper left) shows the results for all segments in this chromosome.

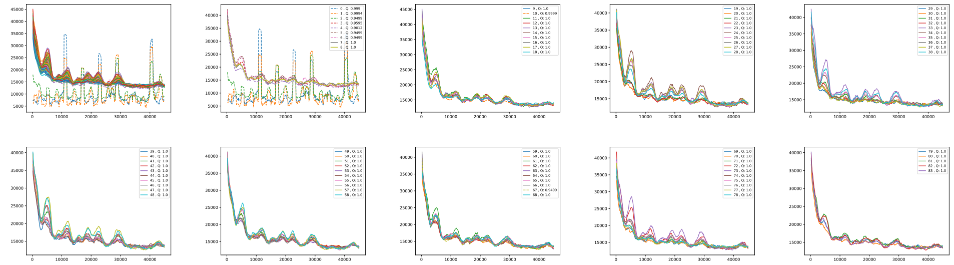

Figure 15: GC/AT content. Fouriers on GC/AT content arrays for chromosome: hg38_chr15. The genome string is divided in adjacent segments of 1Mb; several adjacent segments are covered in each plot, segment numbers are indicated in the legend. First plot (upper left) shows the results for all segments in this chromosome.

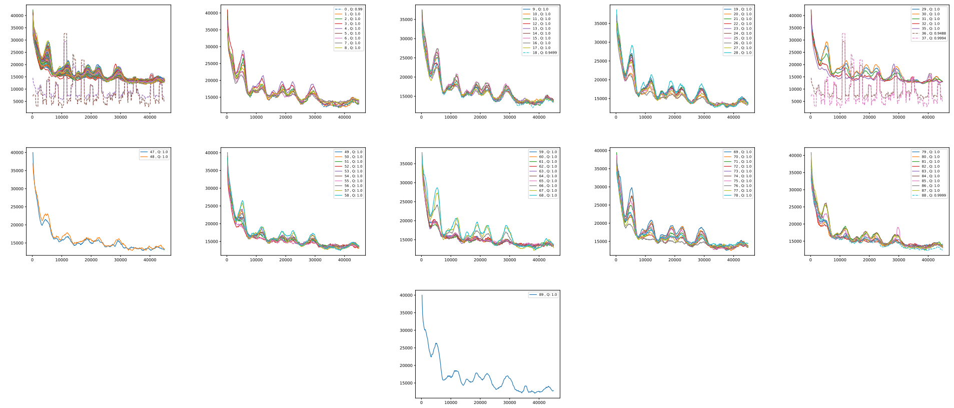

Figure 16: GC/AT content. Fouriers on GC/AT content arrays for chromosome: hg38_chr16. The genome string is divided in adjacent segments of 1Mb; several adjacent segments are covered in each plot, segment numbers are indicated in the legend. First plot (upper left) shows the results for all segments in this chromosome.

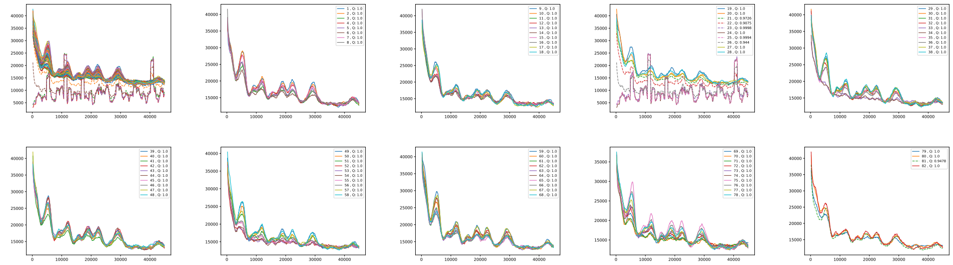

Figure 17: GC/AT content. Fouriers on GC/AT content arrays for chromosome: hg38_chr17. The genome string is divided in adjacent segments of 1Mb; several adjacent segments are covered in each plot, segment numbers are indicated in the legend. First plot (upper left) shows the results for all segments in this chromosome.

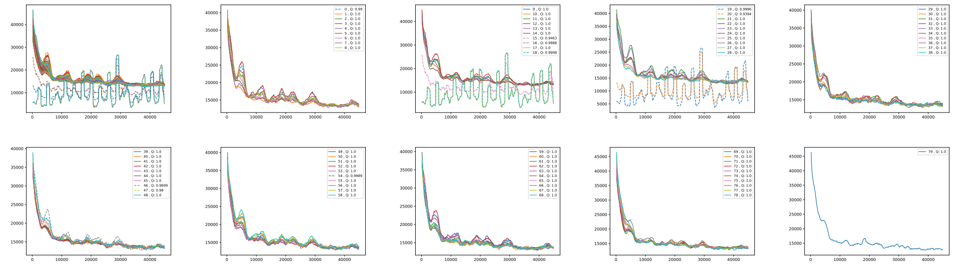

Figure 18: GC/AT content. Fouriers on GC/AT content arrays for chromosome: hg38_chr18. The genome string is divided in adjacent segments of 1Mb; several adjacent segments are covered in each plot, segment numbers are indicated in the legend. First plot (upper left) shows the results for all segments in this chromosome.

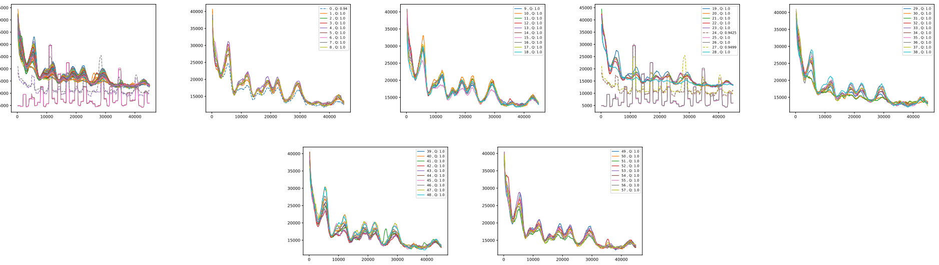

Figure 19: GC/AT content. Fouriers on GC/AT content arrays for chromosome: hg38_chr19. The genome string is divided in adjacent segments of 1Mb; several adjacent segments are covered in each plot, segment numbers are indicated in the legend. First plot (upper left) shows the results for all segments in this chromosome.

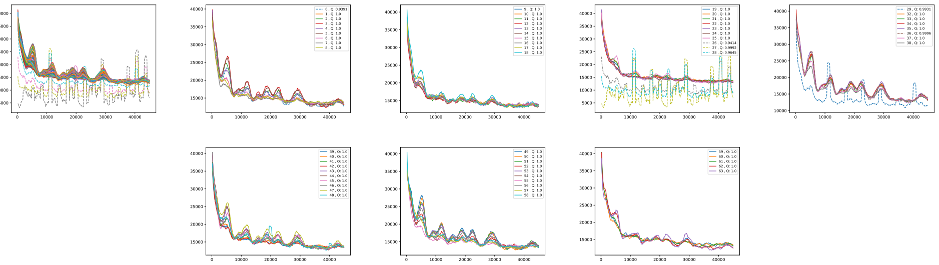

Figure 20: GC/AT content. Fouriers on GC/AT content arrays for chromosome: hg38_chr20. The genome string is divided in adjacent segments of 1Mb; several adjacent segments are covered in each plot, segment numbers are indicated in the legend. First plot (upper left) shows the results for all segments in this chromosome.

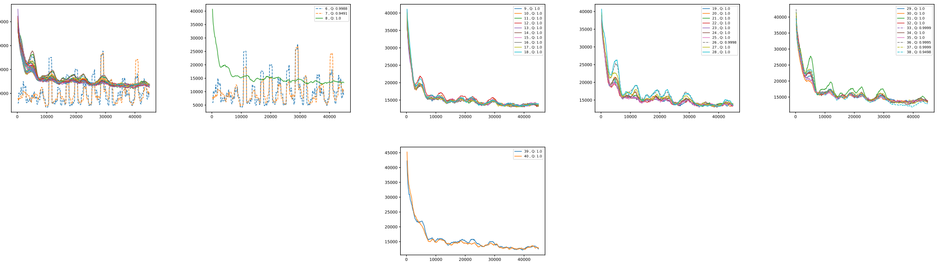

Figure 21: GC/AT content. Fouriers on GC/AT content arrays for chromosome: hg38_chr21. The genome string is divided in adjacent segments of 1Mb; several adjacent segments are covered in each plot, segment numbers are indicated in the legend. First plot (upper left) shows the results for all segments in this chromosome.

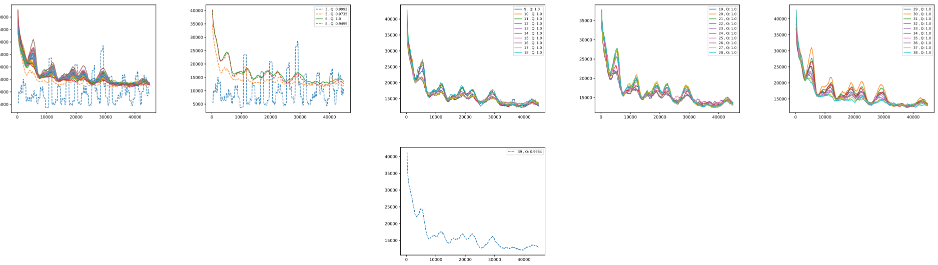

Figure 22: GC/AT content. Fouriers on GC/AT content arrays for chromosome: hg38_chr22. The genome string is divided in adjacent segments of 1Mb; several adjacent segments are covered in each plot, segment numbers are indicated in the legend. First plot (upper left) shows the results for all segments in this chromosome.
