## Supplementary IIe for "Prediction of DNA from context using neural networks"

### Supplementary IIe: Prediction of DNA from context using neural networks. Fouriers, human genome, Markov and $k = 5$ central model (chr22); mouseLSTM4 (chr20).

Christian Grønbæk, Yuhu Liang, Desmond Elliott, and Anders Krogh

November 26, 2020

#### Fouriers plots, predictions on chromosome 22

##### Markov model

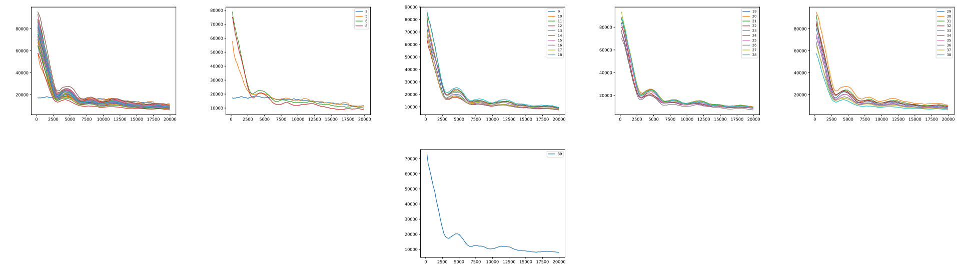

Figure 1: Markov model, k=14. Fouriers on reference-base probability for chromosome: hg38_chr22. The genome string is divided in adjacent segments of 1Mb; several adjacent segments are covered in each plot, segment numbers are indicated in the legend. First plot (upper left) shows the results for all segments in this chromosome.

#### $k = 5$ central model

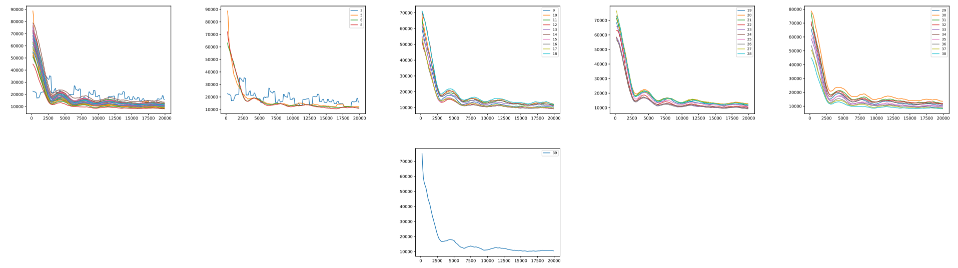

Figure 2: 5-mer model. Fouriers on reference-base probability for chromosome: hg38_chr22. The genome string is divided in adjacent segments of 1Mb; several adjacent segments are covered in each plot, segment numbers are indicated in the legend. First plot (upper left) shows the results for all segments in this chromosome.

#### LSTM1

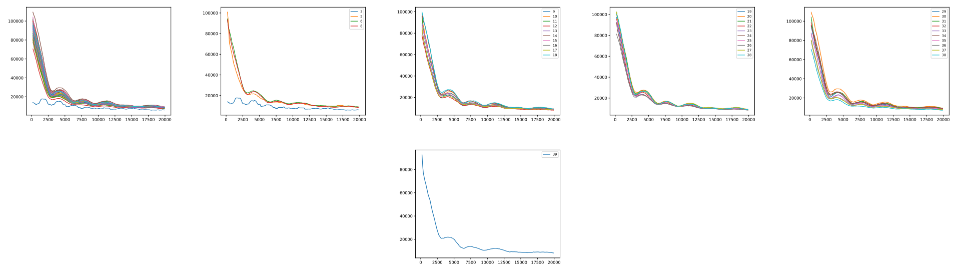

Figure 3: LSTM1. Fouriers on reference-base probability for chromosome: hg38_chr22. The genome string is divided in adjacent segments of 1Mb; several adjacent segments are covered in each plot, segment numbers are indicated in the legend. First plot (upper left) shows the results for all segments in this chromosome.

### Fouriers plots, mouseLSTM4 predictions on chromosome 20

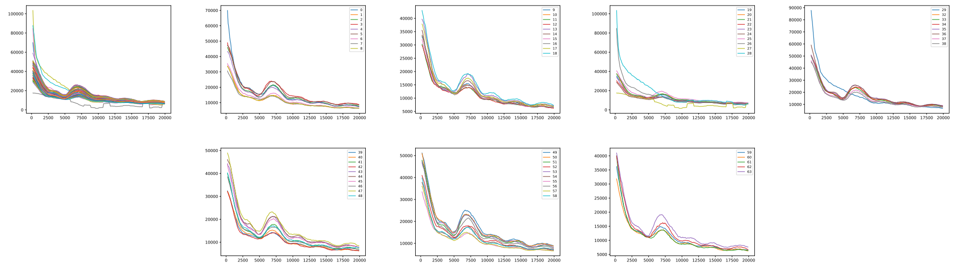

Figure 4: mouseLSTM4. Fouriers on reference-base probability for chromosome: hg38_chr20. The genome string is divided in adjacent segments of 1Mb; several adjacent segments are covered in each plot, segment numbers are indicated in the legend. First plot (upper left) shows the results for all segments in thise chromosome
