## Supplementary VII for "Prediction of DNA from context using neural networks"

22 May 2021

### Single Nucleotide Polymorphisms

In [3] we considered questions related to Single Nucleotide Polymorphisms (SNPs) and established a model of such variants based partially on the Markov model ($k = 14$). Here we make some of the same considerations, but based on the LSTM1 neural network model.

We have tacitly assumed that our models do not depend on the specific genomes on which they were trained, i.e. using genomes of other individuals would give only negligible differences. So we interpret the models as representing an 'evolutionary equilibrium'. Even if the initial occurrence of a SNP may depend on the context dependent nucleotide probabilities, the fixation and frequency in the population is not expected to depend on this. But there could maybe still be some connection: A simple question would be if for (biallelic) disease causing SNPs there is a tendency that the alternative allele is less probable than the reference (given the context) according to the model. Or, if maybe a large difference in model probabilities is indicative of disease causality.

We saw in [3] that none of this seemed to be the case for our simpler models, and despite its superiority it is also not the case for the LSTM1. First though, as shown in Figure 1, we can see how the confidence of LSTM1 carries over to SNPs, here in human chromosome 22 obtained in the 1000Gs Project [1]: The lower right corner, holding SNPs for which the model assigns a probability to the reference base close to 1, is more densely populated for LSTM1 than for the $k = 5$ central model and Markov $k = 14$.

In Figure 2 we consider SNPs from three public databases (1000G again plus ClinVar [2] and COSMIC [4]) on human chromosome 22. The figure shows the density of the difference between the model's probabilities of the reference and the alternative allele for these sets of variants. For comparison 'background' is added: the same density for randomly selected positions on chromosome 22 and for the coding positions among these. Assuming no or very little correlation of SNPs' allele frequencies and the model's probabilities, we should expect such density to be roughly symmetric for an unselected set of SNPs. To see why, consider a SNP having e.g. reference allele A and alternative allele G. By the independence the occurrences of A and G throughout the genome as midpoints in the very same context as at the SNP are independent of the roles of the two bases at the SNP; it is just as likely that G is more frequent as midpoint than A as vice verse. So, in a large, randomly sampled set of SNPs it is equally probable that the probability difference is positive and that it is negative.

As the ClinVar and COSMIC coding variants are disease related, the densities could be non-symmetric; this would be the case if one or both of our questions had an affirmative answer. Now clearly, from the left-hand plot in Figure 2, the densities for both sets are roughly symmetric. So, a tendency of positive difference in model probability is certainly not indicative of a disease causal mutation, nor is a large difference (for the ClinVar set the density seems even to lean towards the left, indicating that for disease causal SNPs the alternative allele is most often the most probable, according to the model). For the cancer related coding variants in the COSMIC set (81163 in total), the density does appear right skewed, but only slightly, and any particular density of SNPs having high difference in probability is absent.

Apparently contrary to our reasoning above, the densities for the 1000Gs SNPs (in total 985834) and the COSMIC non-coding SNPs (196004 in total) on chromosome 22 appear right-skewed having bumps in the tails, much less pronounced in the left-hand tail than in the right (which also follows the background). These bumps turn out to be due to SNPs called in repeats: In the plot to the right in Figure 2 the effect of removing SNPs at repeat positions is shown, and both densities have now become close to symmetric and with no bumps. In [3] we found the same phenomenon for the simpler models. Still, the lack of symmetry needs explanation. One possibility is that the assumed lack of correlation does not hold: in cases of very strongly patterned contexts, e.g. a very clear repeat sequence, a base not following the pattern could very likely be rare both as a variant (across genomes) and as context-midpoint (along a genome). This correlation could very well be coincidental. Another possibility [3] is that these SNPs are artifacts (mapping reads to repeat areas is difficult and can result in wrong calls). Finally, it may show that the reference genome cannot be viewed as an individual genome, because it originates from several individuals, and therefore there are fewer rare SNVs than in an individual genome.

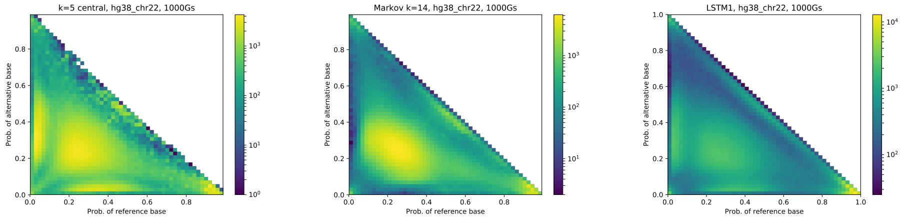


Figure 1: Probability of alternative vs reference base as had by the named models ($k = 5$ central, Markov $k = 14$, LSTM1) for the SNPs in human hg38 chromosome 22 obtained in the 1000Gs Project [1]. Please note that colors are not shared between the plots.

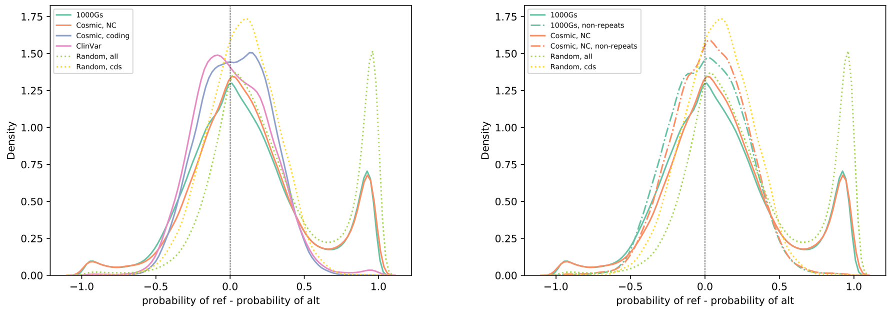


Figure 2: Density of the difference in probability between the reference base and the alternative according to LSTM1 for mutational subsets (of SNPs) on human chromosome 22 as indicated. Left: SNPs as indicated (solid lines) and for a randomly selected set of positions, all or coding positions only (dotted). Right: all SNPs in 1000G and COSMIC non-coding or only those at non-repeat positions; backgrounds as in plot to the left.
